# *Ppd-1* Remodels Spike Architecture by Regulating Floral Development in wheat

**DOI:** 10.1101/2020.05.11.087809

**Authors:** Yangyang Liu, Lili Zhang, Michael Melzer, Liping Shen, Zhiwen Sun, Ziying Wang, Thorsten Schnurbusch, Zifeng Guo

## Abstract

The determination of spike architecture is critical to grain yield in wheat (*Triticum aestivum*), yet the underlying mechanisms remain largely unknown. Here, we measured 51 traits associated with spike architecture and floral development in 197 wheat accessions with photoperiod sensitive and insensitive alleles. We included five distinct allele combinations at the *Photoperiod-1* (*Ppd-1*) loci. A systematic dissection of all recorded phenotypes revealed connections between floral development, spike architecture and grain yield. Modifying the durations of spikelet primordia initiation did not necessarily affect spikelet number. In addition, *Ppd-1* loci clearly influenced rachis dry weight, pointing to the rachis vascular system as a potential target for higher yield. *Ppd-1* displayed opposite effects on the durations of pre and post-anthesis phases. *Ppd-1* controlled carpel size, but not anther size. Finally, the photoperiod-insensitive alleles of *Ppd-1* triggered floral degeneration. In parallel, we profiled the spike transcriptome at six stages and four positions in three *Ppd-1* genotypes which consists of 234 samples. Integrating phenotypic and expression data suggested that loss of function in *Ppd-1* loci delayed floral degeneration by regulating autophagy and extended floret development by regulating genes in different families. Therefore, we concluded that *Ppd-1* remodels spike architecture by regulating floral development in wheat.

## Introduction

Wheat (*Triticum aestivum* L.) is one of the top three crops cultivated worldwide, based on tons produced each year and arable land dedicated to its cultivation. Improving grain yield is a crucial target in wheat breeding, of which grain number is a key factor, itself determined by spike architecture. A spike generally consists of 20-30 spikelets arranged along the main inflorescence stem (the rachis). Each spikelet may produce 8-12 floret primordia (potential grains) in hexaploid wheat (Guo and Schnurbusch, 2015). However, most floret primordia degenerate during the pre-anthesis stage, leaving 0-4 fertile floret per spikelet, drastically reducing potential grain yield. Floral development and abortion during pre-anthesis may be divided into seven distinct stages on the basis of floret morphology: terminal spikelet (TS), white anther (WA), green anther (GA), yellow anther (YA), tipping (TP), heading (HD), and anthesis (AN). The main feature of TS stage is the completion (or termination) of spikelet initiation (Kirby and Appleyard, 1987). At this stage, the last few primordia do not develop into spikelets but instead form the glumes and floret primordia of a terminal spikelet(Kirby and Appleyard, 1987). During WA stage, the meristematic dome initiates 7-9 floret primordia within each individual spikelet (1-3 primordia fewer than the maximum number)(Kirby and Appleyard, 1987). The lemmas of florets 1 and 2 at the base of the spikelet completely enclose the anther and ovary(Kirby and Appleyard, 1987). At the GA stage, the meristematic dome completes the initiation of floret primordia, so that after this stage no more floret primordia will be produced (Guo and Schnurbusch, 2015); the glumes almost completely cover all but the tips of the florets. The spike length peaks at the YA stage (Guo et al., 2018); glumes are fully formed, and the lemmas of the first three florets are visible. At the TP stage, the first awn is visible (Zadoks et al., 1974), at HD stage, half of the spike emerges from the last leaf blade (Zadoks et al., 1974). Finally, at the AN stage, yellow anthers appear along the spike (Zadoks et al., 1974).

Spike architecture is a complicated trait, that is determined by multiple factors. Spike fertility index (i.e. the ratio between grain number per spike and weight of spike chaff) is a critical metric of assimilate distribution between grains and spike chaff (e.g. glume, lemma, palea, rachis). Grain number per spikelet along the main inflorescence also influences spike shape. Spikelet density (the ratio between spike length and spikelet number) also indicates spike compactness. Spikelet fertility (the ratio between fertile and total spikelet number) informs on the proportion of aborted spikelets. In addition, grain size traits (thousand-kernel weight-TKW, grain area, grain width, grain length) all contribute to spike architecture.

The response to photoperiod is major agronomic trait that modulates flowering time. In wheat, photoperiod sensitivity is controlled by three *Photoperiod-1* (*Ppd-1*) genes that are located on chromosomes 2A, 2B, and 2D and designated *Ppd-A1, Ppd-B1*, and *Ppd-D1*, respectively (Scarth and Law, 1983; Wilhelm et al., 2009). *Ppd-1* genes belong to the PSEUDO-RESPONSE REGULATOR family of circadian clock and photoperiod regulators. Their photoperiod insensitive alleles are designated by the suffix “a” (e.g., *Ppd-1a*), and alleles conferring a photoperiod sensitivity have the suffix “b” (e.g., *Ppd1b*) (Mclntosh et al., 2003; Shaw et al., 2013). In wheat breeding, photoperiod insensitivity has been widely introduced in cultivars where early flowering is desirable to avoid stress or to allow crops to flower even under short-day photoperiods (Kato and Yokoyama, 1992; Worland et al., 1998; Shaw et al., 2013). *Ppd-1b* alleles follow a diurnal expression pattern: they are highly expressed during the light period, whereas their expression falls to very low levels in dark. By contrast, *Ppd-1a* alleles lack this diurnal pattern, displaying a constant and high expression pattern (Beales et al., 2007;Wilhelm et al., 2009). *Ppd-1a* alleles raise the expression of the wheat *Flowering Locus 1* (*TaFT1*) in leaves, from which the FT protein later moves to the shoot apex to induce flowering (Shaw et al., 2012). *Ppd-1* was previously reported to similarly play a key role during supernumerary spikelet formation in wheat (Boden et al., 2015).

Here, we first determined the effects of *Ppd-1* loci on spike architecture and developmental traits in a natural population of 197 wheat accessions, then contributions of *Pp*d*-1* alleles to the measured traits using five *Ppd-1* genotypes that vary in their observed flowering time. By quantifying changes in the transcriptome at six stages and four positions along the spike and in three *Ppd-1* genotypes, we demonstrate that *Ppd-1* remodels spike architecture by regulating floral development in wheat.

## Results

### *Ppd-1* loci affect spike architecture, floret development, and grain yield in 197 wheat accessions

To determine the effects of the *Ppd-D1* locus on spike architecture and floret development, we genotyped the *Ppd-D1* gene in a wheat population of 197 wheat accessions and phenotyped them for 51 traits related to spike architecture and development. We then calculated spike architecture traits separately for photoperiod-sensitive (*Ppd-D1b*) and photoperiod-insensitive (*Ppd-D1a*) allelic variants at the *Ppd-D1* locus (Table 1).

**Table 1.** Effects of sensitivity and insensitivity alleles at the *Ppd-D1* (chromosome 2D) locus on spike architecture and floret development traits.

| Traits | Sensitivity (n=165) | Insensitivity (n=32) | Difference (%) | T-test |
| --- | --- | --- | --- | --- |
| Final Spike length (cm) | 11.47 ± 1.29 | 9.71 ± 1.07 | -15 | 1.53E-11*** |
| Total spikelet number | 25.28 ± 2.87 | 22.03 ± 2.1 | -13 | 6.71E-09*** |
| Fertile spikelet number | 22.21 ± 2.81 | 20.28 ± 1.81 | -9 | 0.0003*** |
| Spikelet fertility | 0.89 ± 0.09 | 0.92 ± 0.04 | +4 | 0.0248* |
| Spike DW (g) | 3.76 ± 0.68 | 3.56 ± 0.4 | -5 | 0.1115 |
| Grain number per spike | 68.75 ± 12.58 | 61.82 ± 8.63 | -10 | 0.0092** |
| Grain weight per spike spike | 2.78 ± 0.6 | 2.73 ± 0.31 | -2 | 0.7177 |
| Spike chaff (g) | 1.31 ± 0.95 | 1.43 ± 1.21 | +9 | 0.5575 |
| TKW (g) | 40.87 ± 6.74 | 44.9 ± 6.42 | +10 | 0.0063** |
| Spikelet density | 2.22 ± 0.24 | 2.28 ± 0.21 | +3 | 0.1664 |
| Spike fertility index | 72.03 ± 14.5 | 79 ± 14.52 | +10 | 0.0288* |
| Grain weight: spike chaff | 2.91 ± 0.64 | 3.51 ± 0.61 | +20 | 2.60E-05*** |
| Max.Floret A | 9.1 ± 0.55 | 8.83 ± 0.48 | -3 | 0.0151* |
| Max.Floret C | 9.87 ± 0.59 | 9.34 ± 0.51 | -5 | 6.20E-06*** |
| Max.Floret B | 9.72 ± 0.7 | 9.13 ± 0.56 | -6 | 2.15E-05*** |
| Grain A | 2.6 ± 0.77 | 2.97 ± 0.52 | +14 | 0.011* |
| Grain C | 3.71 ± 0.5 | 3.85 ± 0.39 | +4 | 0.127 |
| Grain B | 3.56 ± 0.6 | 3.57 ± 0.45 | +0.3 | 0.9233 |
| Survival A | 0.29 ± 0.09 | 0.33 ± 0.06 | +16 | 0.0051** |
| Survival C | 0.38 ± 0.05 | 0.41 ± 0.04 | +9 | 0.0013** |
| Survival B | 0.37 ± 0.07 | 0.39 ± 0.05 | +5 | 0.1177 |
| Floret primordia loss A | 6.5 ± 0.96 | 5.59 ± 1.86 | -14 | 8.48E-05*** |
| Floret primordia loss C | 6.15 ± 0.76 | 5.19 ± 1.78 | -16 | 1.95E-06*** |
| Floret primordia loss B | 6.15 ± 0.93 | 5.28 ± 1.85 | -14 | 0.0001*** |
| TS spike length (cm) | 0.37 ± 0.07 | 0.39 ± 0.08 | +5 | 0.1861 |
| WA spike length (cm) | 2.08 ± 0.4 | 2.28 ± 0.4 | +9 | 0.0225* |
| GA spike length (cm) | 5.52 ± 0.91 | 5.18 ± 0.77 | -6 | 0.0591 |
| YA spike length (cm) | 10.49 ± 1.28 | 9.32 ± 1.16 | -11 | 6.01E-06*** |
| TP spike length (cm) | 11.94 ± 1.28 | 9.97 ± 1.03 | -17 | 1.73E-13*** |
| HD spike length (cm) | 12.09 ± 1.28 | 10.13 ± 1.1 | -16 | 5.21E-13*** |
| AN spike length (cm) | 11.99 ± 1.32 | 10.1 ± 1.19 | -16 | 8.85E-12*** |
| TS stage (thermal time °Cd) | 1035.25 ± 101.57 | 888.81 ± 78.02 | -14 | 8.25E-13*** |
| WA stage (thermal time °Cd) | 1239.52 ± 99.38 | 1057.52 ± 99.1 | -15 | 2.10E-16*** |
| GA stage (thermal time °Cd) | 1328.57 ± 105.4 | 1130.88 ± 110.18 | -15 | 5.78E-18*** |
| YA stage (thermal time °Cd) | 1398.72 ± 117.69 | 1187.97 ± 112.76 | -15 | 4.34E-17*** |
| TP stage (thermal time °Cd) | 1468.08 ± 136.22 | 1235.88 ± 112.17 | -16 | 2.19E-16*** |
| HD stage (thermal time °Cd) | 1527.87 ± 144.92 | 1282 ± 109.21 | -16 | 1.73E-16*** |
| AN stage (thermal time °Cd) | 1645.19 ± 153.89 | 1367 ± 118.79 | -17 | 1.04E-17*** |
| Grain yield (ENV 1, T/ha) | 99.01 ± 6.23 | 98.07 ± 4.31 | -1 | 0.4138 |
| Grain yield (ENV 2, T/ha) | 106.88 ± 6.13 | 101.84 ± 6.69 | -5 | 4.61E-05*** |
| Grain yield (ENV 3, T/ha) | 92.69 ± 6.35 | 95.35 ± 5.15 | +3 | 0.0275* |
| Grain yield (ENV 4, T/ha) | 100.19 ± 5.14 | 99.95 ± 4.75 | -0.2 | 0.8038 |
| Grain yield (ENV 5, T/ha) | 91.70 ± 5.49 | 94.08 ± 4.17 | +3 | 0.0216* |
| Grain yield (ENV 6, T/ha) | 104.06 ± 5.79 | 103.99 ± 4.44 | -0.1 | 0.9537 |
| Grain yield (ENV 7, T/ha) | 79.93 ± 6.41 | 76.04 ± 6.86 | -5 | 0.0023** |
| Grain yield (ENV 8, T/ha) | 110.98 ± 5.96 | 107.27 ± 6.38 | -3 | 0.0018** |
| Spike DW (ENV 3, g) | 32.19 ± 3.52 | 31.22 ± 3.75 | -3 | 0.1622 |
| Spike DW (ENV 8, g) | 33.14 ± 3.72 | 32.97 ± 3.51 | -1 | 0.8167 |
| Grain number (ENV 1) | 51.23 ± 7.20 | 47.54 ± 8.42 | -7 | 0.0112* |
| Grain number (ENV 3) | 54.87 ± 6.47 | 49.60 ± 7.06 | -10 | 5.39E-05*** |
| Grain number (ENV 4) | 47.60 ± 5.27 | 43.81 ± 5.18 | -8 | 2.60E-04** |

The spike length was measured without the awn. The spike chaff refers to the rachis with empty spikelets after removal of grains. Spike fertility index, spikelet density, and fertility were determined based on spikelet number per cm of spike length, grain number (per spike) g<sup>-1</sup> of spike chaff (per spike) at harvest, and the ratio of the fertile and total spikelet numbers. Infertile spikelets were defined as spikelets that did not set any grain (i.e. were completely empty), Fertile spikelets produced at least one grain. Grain A/Max.Floret A, grain C/Max.Floret C, grain B/Max.Floret B refer to the grain number per spikelet/maximum number of floret primordia per spikelet at apical (the third spikelet from the top of spike), central (the spikelet in the center of spike), basal (the third spikelet from the bottom of spike) spikelet, respectively. Survival/floret primordia loss are the ratios/differences between grain number and maximum number of floret primordia. The data were presented as the mean±SD (difference %, P value for T test). The percentage of differences between NIL/KL and WT were are given relative to the values of sensitive alleles. \*, \*\*, and \*\*\* indicate significance levels of 0.05, 0.01, and 0.001, respectively. For floret development traits: n=3; for the spike architecture traits: n=6.

The allelic status at the *Ppd-D1* locus significantly affected 37 traits. When normalized to phenotypic values measured for the photoperiod sensitive *Ppd-D1b* alleles, variation ranged from 3 to 20% (when *Ppd-D1a* alleles have a positive effect on the trait) and from 3 to 17% (when *Ppd-D1a* alleles negatively affected a trait). Photoperiod-insensitive *Ppd-D1a* alleles affected

spike compactness by lowering final spike length by 15%, while also decreasing total spikelet number by 13% and the number of fertile spikelets by 9%. Because of this unequal effect on total and fertile spikelet numbers, *Ppd-D1a* alleles resulted in a modest increase in spikelet fertility (the ratio between fertile and total spikelet number) of 4%, although the statistical significance was weak. However, we did not observe a significant difference in spikelet density (the ratio between total spikelet number and spike length). In addition, *Ppd-D1a* (the insensitive allele) increased TKW, spike fertility index (that is the ratio between grain number per spike and weight of spike chaff) and the ratio of grain weight/spike chaff by 10%, 10% and 20%, respectively, compared with *Ppd-D1b* (the sensitive allele).

One measure of relative spikelet fertility captures the maximum number of floret primordia in three representative spikelets: the apical (A) and basal (B) florets of the third spikelet from the top or bottom of the spike, respectively, while the central floret (C) sits at the center of the spike. *Ppd-D1a* insensitive alleles resulted in small drop in the maximum number of floret primordia per spikelet at each A, C and B position (annotated as Max. Floret A/C/B in Table 1): 3% for A spikes, 5% for C spikes and 6% for B spikes, compared with the photoperiod-sensitive allele. Looking at the number of grains produced by the apical spikelet, *Ppd-D1a* insensitive alleles increased the grain number per spikelet (grain A) by 14% and floret survival by 16% (calculated as the ratio between maximum number of floret primordia and grain number, and labeled survival A in Table 1). Moreover, *Ppd-D1a* insensitive alleles led to a marked reduction of the loss of floret primordia (expressed as the difference between maximum number of floret primordia and grain number) by 14% (floret primordia A and B) and 16% (floret primordia C). Moreover, *Ppd-D1a* insensitive alleles reduced spike dry weight (DW) by 5% and the length of all spike developmental stages by 14–17%. That *Ppd-D1a* insensitive alleles lowered maximum number of floret primordia, spike length and spike DW may be attributable to their shortening of each spike developmental stage (Table 1).

We also determined grain yield in eight different field environments, as well as spike dry weight in two field environments and grain number in three field environments. *Ppd-D1b* sensitive alleles consistently increased spike dry weight and grain number, possibly as a consequence of their prolonging spike developmental stages. Since grain yield is a complicated trait that may be influenced by tiller number, grain number, and TKW, *Ppd-D1a* insensitive alleles had significant positive effects on grain yield in two environments of ∼3% and significant negative effects in three environments of 3–5%. These results show that allelic variants of *Ppd-D1* clearly influenced spike architecture and floret developmental traits.

### Validating the effects of *Ppd-1* loci on spike architecture and floral development traits in *Ppd-1* mutants

To further validate the effects of *Ppd-1* loci on spike architecture and development, we measured the same spike- and spikelet-associated phenotypes in five strains differing at the *Ppd-1* loci (see plant materials). The spring wheat variety Paragon carries photoperiod-sensitive alleles in all three genomes: *Ppd-A1bB1bD1b*, and displays an intermediate flowering time characteristic of this wild type cultivar. Two near-isogenic lines (NILs) have only *Ppd-1* insensitive alleles (*Ppd-A1aB1aD1a*) that result in early flowering. We also characterized two independent mutant lines with loss of function alleles at all three loci (*ppd-A1 B1 D1*) that confer late flowering. Although the two NILs obtained their insensitive alleles from different genotypes, they displayed similar spike architecture and floret developmental traits. Similarly, the mutant alleles combined into the two mutant lines originated from different germplasm but showed similar phenotypes. We therefore determined the average phenotypic values for *Ppd-A1aB1aD1a* and *ppd-A1B1D1* by calculating the average of both NILs and both mutant lines.

#### Ppd-1 also affects floral development and abortion

As expected, *Ppd-1* insensitive alleles reduced the time required to complete each of the six stages by 12–17% when compared to the wild type (WT, carrying only sensitive alleles). By contrast, *Ppd-1* loss of function alleles increased the length of each stage by 19–34% (Table 2). The duration of each stage required for spikelet and floret development and abortion events therefore varied among the three genotypes according to their *Ppd-1* genotypes. The timing of visible floral primordium termination similarly changed as a function of the *Ppd-1* genotype, as shown in Table 2. Indeed, visible floral degradation started during the YA stage and extended into the TP stage in early-flowering NILs (carrying only insensitive alleles, *Ppd-A1aB1aD1a*). The abortion of apical florets occurred slightly later in both WT and mutant lines, starting during the TP stage and ending with the HD stage. These results therefore indicated that *Ppd-1* insensitive alleles may trigger floral degeneration.

**Table 2.** Numbers of floret primordia at seven stages in WT, NIL and KL.

| Genotype | TS | WA | GA | YA | TP | HD | AN |
| --- | --- | --- | --- | --- | --- | --- | --- |
| NIL | 5.67±0.47 | 8.80±0.40 | 10.40±0.49 | 10.67±0.75<sup>1</sup> | 5.60±0.49<sup>1</sup> | 5.57±0.49 | 5.33±0.47 |
| WT | 6.40±0.49 | 8.82±0.83 | 10.70±0.64 | 10.38±0.48 | 9.80±1.08<sup>1</sup> | 6.00±0.53<sup>1</sup> | 5.70±0.46 |
| KL | 5.00±0.82 | 8.91±0.67 | 11.63±0.48 | 11.29±0.70 | 11.00±0.63<sup>1</sup> | 5.80±0.40<sup>1</sup> | 5.57±0.49 |

The *Ppd-1* genotypes for the three lines are *Ppd-A1bB1bD1b* for wild type (WT), *Ppd-A1aB1aD1a* for the near-isogenic line (NIL) and *ppd-A1B1D1* for the triple mutant line (KL). *Ppd-A1bB1bD1b* are *Ppd-1* sensitive (late flowering) alleles in the A, B and D genomes; *Ppd-A1aB1aD1a* are *Ppd-1*-insensitive (early flowering) alleles in the A, B and D genomes; and *ppd-A1*, *ppd-B1* and *ppd-D1* represent loss of function alleles of *Ppd-1* in the A, B and D genomes, as the flowering times of plants with these genotypes were later than that of the wild type. The changes in the numbers of floret primordia within individual spikelets were determined across the 7 spike developmental stages: terminal spikelet stage (TS, the stage before the WA stage), WA, GA, YA, TP, HD and AN. At the TS stage, there is no obvious floral structure (e.g., carpel and anther), so we collected samples for transcriptome from tipping stage. Data are presented as the mean ± SD, n=3 for all the stage. <sup>1</sup> Indicates apical floral abortion.

We observed the maximum number of floret primordia at the GA stage in all three *Ppd-1* genotypes (Table 2), although the genotype did influence this number as well, as the maximum number of floret primordia was higher in mutant lines compared to NILs and WT. This observation suggests that an extended time window of floral initiation (from the TS to GA stage) in mutant lines increased the number of floret primordia. Although both WT and mutant lines took longer to complete each developmental stage, this did not result in a significant increase in fertile florets per spikelet, as measured at the AN stage, relative to NILs.

#### Ppd-1 markedly affects ovary width but not anther length

We determined carpel width and anther length during the AN stage in three *Ppd-1* genotypes NILs (early-flowering, *Ppd-A1aB1aD1a*), WT (intermediate flowering, *Ppd-A1bB1bD1b*), and mutant lines (late flowering, *ppd-A1B1D1*) in the first 4 florets starting from the base of the spike, designated F1 to F4 (Table 3). The mutant lines displayed increased carpel width at all 4 floret positions compared to WT in the range of 9-19%, while the NILs showed reduced carpel width by 12-26% (Table 3). We hypothesize that the extension of the floret growth period by 67% (between the TS and AN stages) in the mutant lines relative to WT allowed more time for carpel growth and led to wider carpels. Conversely, the 8% shorter time for floret growth experienced by the NILs resulted in thinner carpels over the same developmental window. In addition, the genotypes at the *Ppd-1* loci affected the lengths of all six stages equally: photoperiod-insensitive alleles accelerated all stages by 12-18%, and *ppd-1* loss of function alleles delayed each stage by 27-34%. By contrast, anther length remained relatively stable among the three *Ppd-1* genotypes.

**Table 3.**
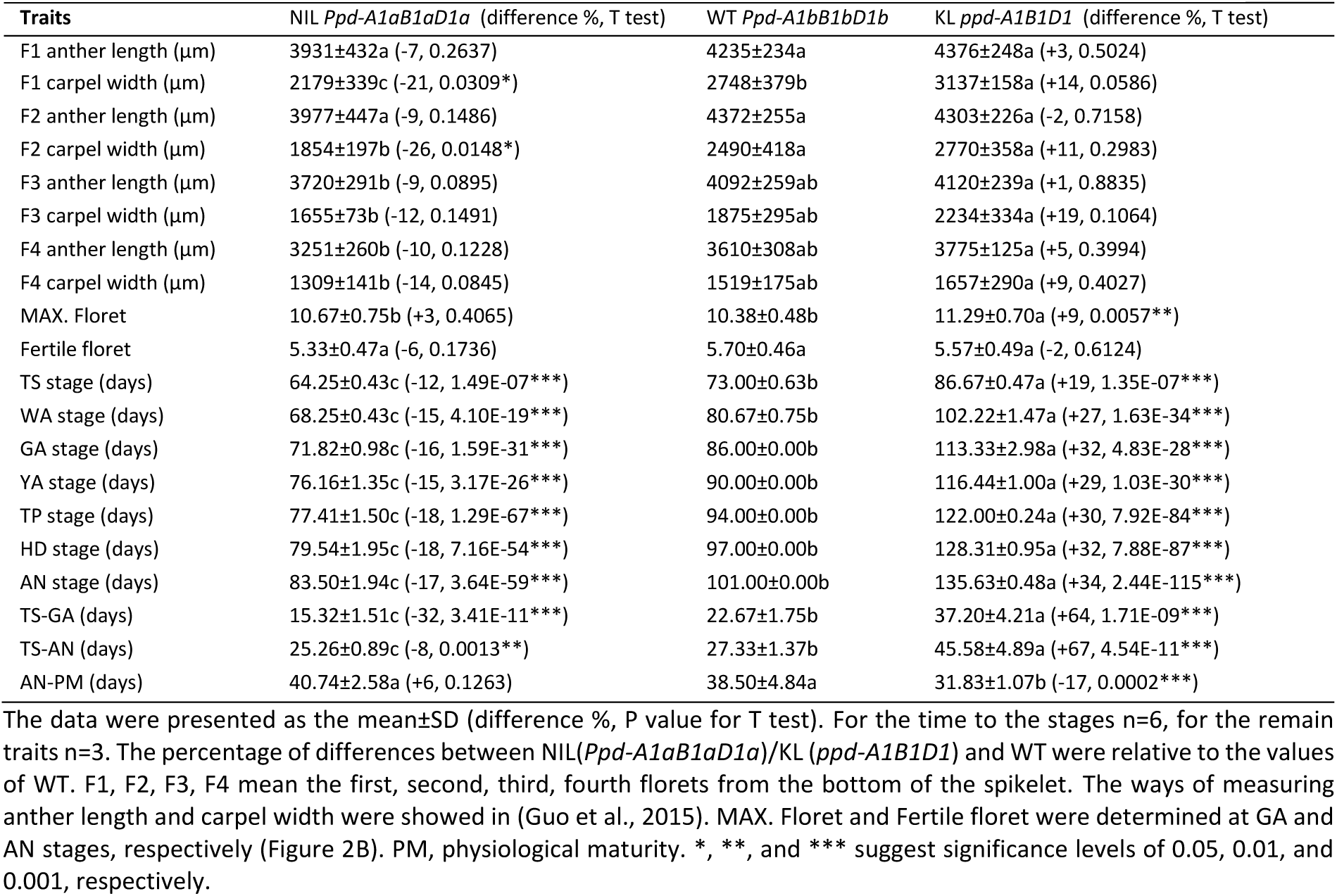
Effects of *Ppd-1* on floret primordia initiation and development traits.

| Traits | NIL *Ppd-A1aB1aD1a* (difference %, T test) | WT *Ppd-A1bB1bD1b* | KL *ppd-A1B1D1* (difference %, T test) |
| --- | --- | --- | --- |
| F1 anther length (μm) | 3931±432a (-7, 0.2637) | 4235±234a | 4376±248a (+3, 0.5024) |
| F1 carpel width (μm) | 2179±339c (-21, 0.0309*) | 2748±379b | 3137±158a (+14, 0.0586) |
| F2 anther length (μm) | 3977±447a (-9, 0.1486) | 4372±255a | 4303±226a (-2, 0.7158) |
| F2 carpel width (μm) | 1854±197b (-26, 0.0148*) | 2490±418a | 2770±358a (+11, 0.2983) |
| F3 anther length (μm) | 3720±291b (-9, 0.0895) | 4092±259ab | 4120±239a (+1, 0.8835) |
| F3 carpel width (μm) | 1655±73b (-12, 0.1491) | 1875±295ab | 2234±334a (+19, 0.1064) |
| F4 anther length (μm) | 3251±260b (-10, 0.1228) | 3610±308ab | 3775±125a (+5, 0.3994) |
| F4 carpel width (μm) | 1309±141b (-14, 0.0845) | 1519±175ab | 1657±290a (+9, 0.4027) |
| MAX. Floret | 10.67±0.75b (+3, 0.4065) | 10.38±0.48b | 11.29±0.70a (+9, 0.0057**) |
| Fertile floret | 5.33±0.47a (-6, 0.1736) | 5.70±0.46a | 5.57±0.49a (-2, 0.6124) |
| TS stage (days) | 64.25±0.43c (-12, 1.49E-07***) | 73.00±0.63b | 86.67±0.47a (+19, 1.35E-07***) |
| WA stage (days) | 68.25±0.43c (-15, 4.10E-19***) | 80.67±0.75b | 102.22±1.47a (+27, 1.63E-34***) |
| GA stage (days) | 71.82±0.98c (-16, 1.59E-31***) | 86.00±0.00b | 113.33±2.98a (+32, 4.83E-28***) |
| YA stage (days) | 76.16±1.35c (-15, 3.17E-26***) | 90.00±0.00b | 116.44±1.00a (+29, 1.03E-30***) |
| TP stage (days) | 77.41±1.50c (-18, 1.29E-67***) | 94.00±0.00b | 122.00±0.24a (+30, 7.92E-84***) |
| HD stage (days) | 79.54±1.95c (-18, 7.16E-54***) | 97.00±0.00b | 128.31±0.95a (+32, 7.88E-87***) |
| AN stage (days) | 83.50±1.94c (-17, 3.64E-59***) | 101.00±0.00b | 135.63±0.48a (+34, 2.44E-115***) |
| TS-GA (days) | 15.32±1.51c (-32, 3.41E-11***) | 22.67±1.75b | 37.20±4.21a (+64, 1.71E-09***) |
| TS-AN (days) | 25.26±0.89c (-8, 0.0013**) | 27.33±1.37b | 45.58±4.89a (+67, 4.54E-11***) |
| AN-PM (days) | 40.74±2.58a (+6, 0.1263) | 38.50±4.84a | 31.83±1.07b (-17, 0.0002***) |

### Differences of Floral Development Induced by *Ppd-1* Loci Explain their Effects on Spike Architecture

When compared to WT (with photoperiod-sensitive *Ppd-1* alleles and intermediate flowering time), the photoperiod-insensitive *Ppd-1* alleles in NILs greatly reduced a number of spike-related traits: spike length (24% decrease), total spikelet number (26% decrease), and fertile spikelet number (24% decrease) (Table 4). We hypothesized that the reduction in the duration of spikelet primordia initiation caused by the loss of function *ppd-1* alleles carried by the mutant lines (and responsible for late flowering) during the TS stage was responsible for these negative effects (Table 3). However, the clear extension of spikelet primordia production during the TS stage was not accompanied by a significant change in spikelet number. The photoperiod-insensitive *Ppd-1* alleles in NILs positively influenced spikelet fertility (the ratio between fertile spikelet number and total spikelet number), suggesting that lower total spikelet numbers caused a less fraction of spikelets to abort (Table 4, Figure 1).

**Table 4.**
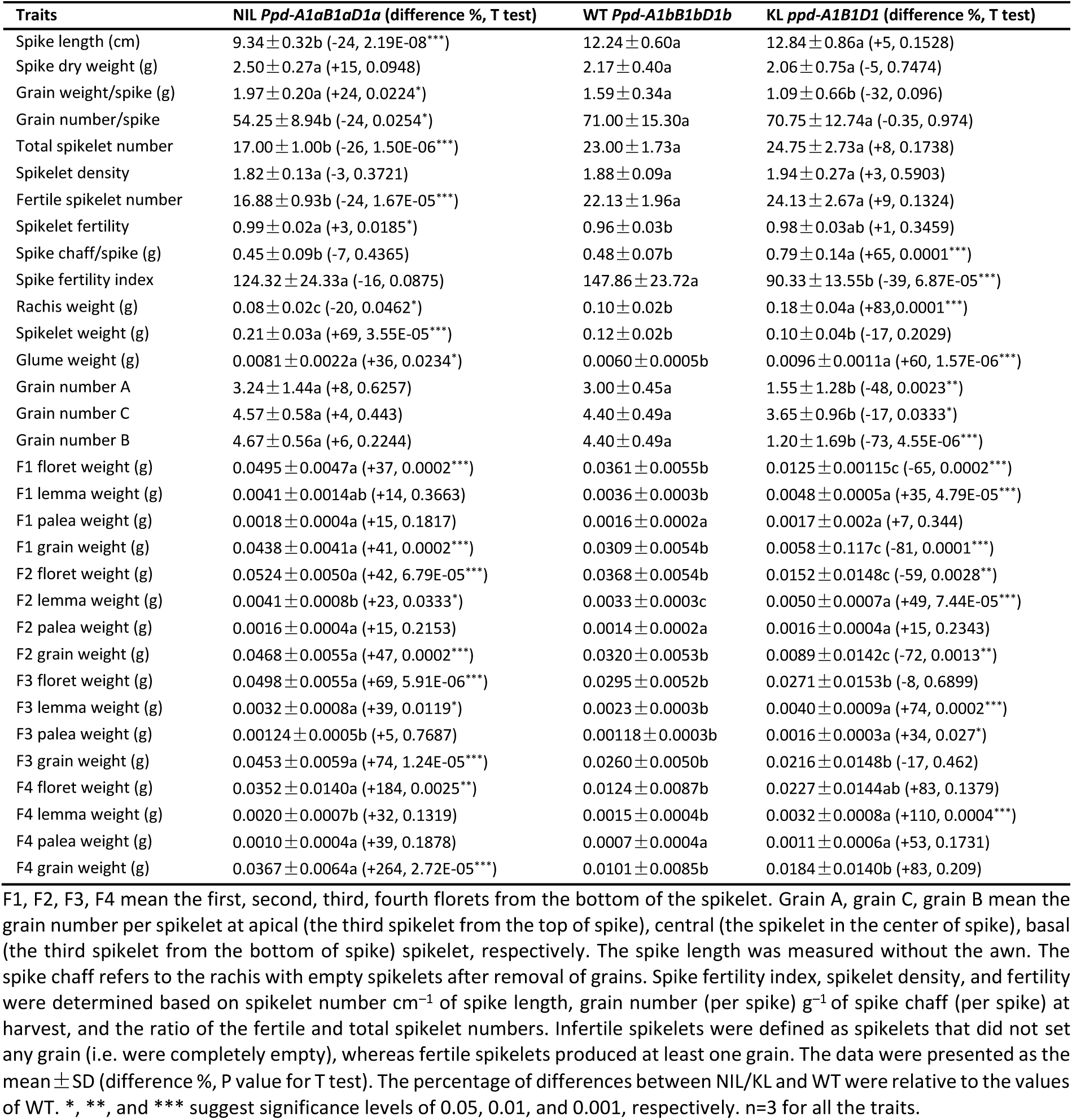
Effects of *Ppd-1* genotypes on spike architecture and floral development traits.

**Figure 1.**
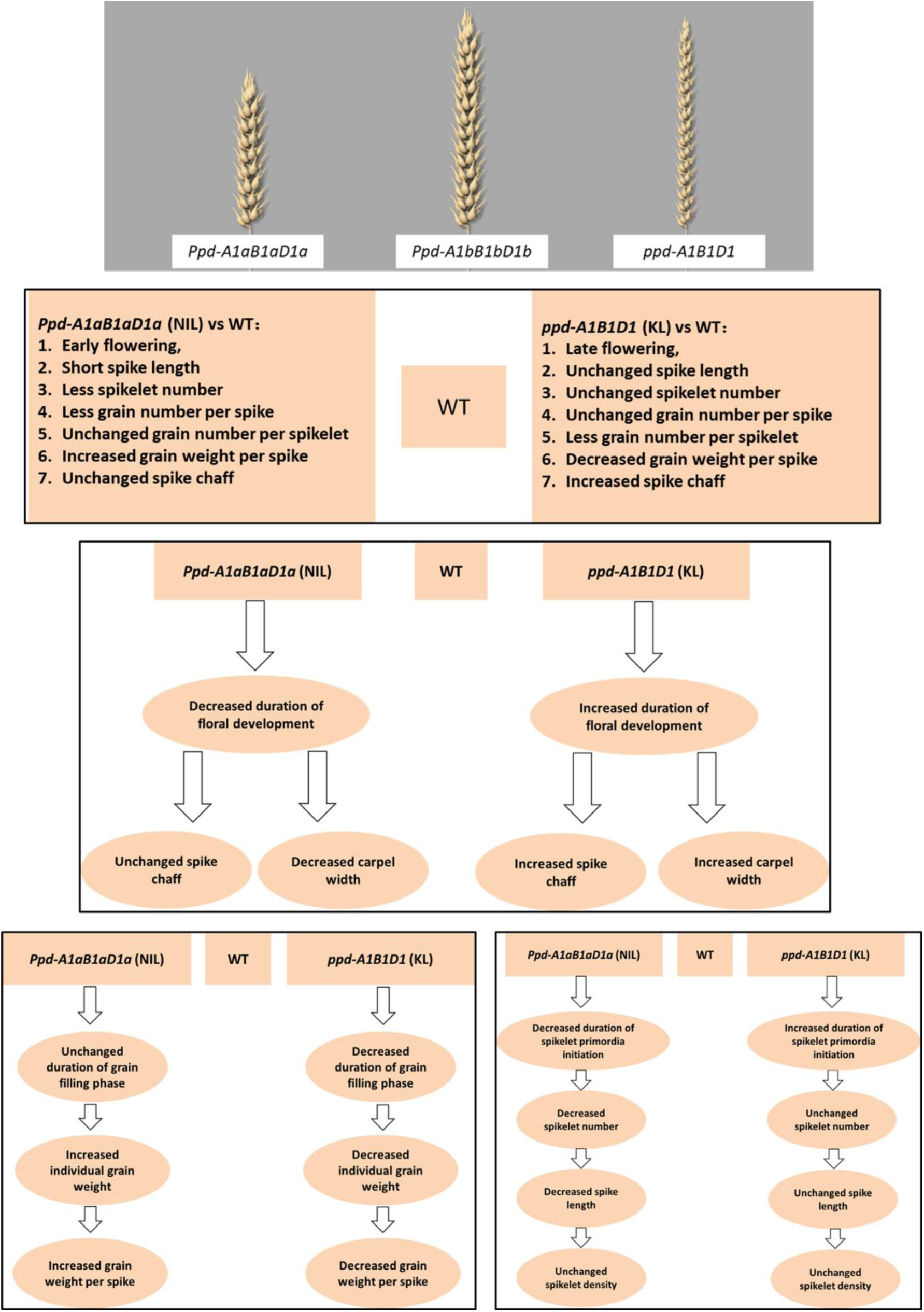
Modification of spike architecture and floral development traits as a function of *Ppd-1* genotypes. Phenotypic differences between NILs (carrying only *Ppd-1* insensitive alleles, *Ppd-A1aB1aD1a*) and mutant lines (loss of function alleles, *ppd-A1B1D1*) compared with WT (*Ppd-1* sensitive alleles, *Ppd-A1bB1bD1b*) in wheat. Unchanged: no significant difference between *Ppd-1* genotypes.

Surprisingly, we observed opposite effects on grain number per spikelet and grain number per spike caused by the photoperiod-insensitive *Ppd-1* alleles in NILs: they reduced grain number per spike in NILs relative to WT by 24%, while raising grain number per spikelet (relative to WT) by 8% for grain A, 4% for grain C, and 6% for grain B (Table 4, Figure 1). The mutant lines exhibited a significant increase in fertile spikelet number, which may be explained by the opposite effects on grain number/spikelet/spike brought on by *ppd-1* loss of function alleles (Table 4, Figure 1). Compared to WT, NILs increased grain weight per spike by 24%. Consistent with the proposed roles of *Ppd-1* alleles, mutant lines lowered the same phenotype by 32%.

The *ppd*-*1* alleles in the mutant lines significantly increased spike chaff weight (the dry weight of a spike after removal of grains) by 65% (Table 4, Figure 1), which may be attributed to their lengthening of floret development stages (TS-AN, +67%, Table 3). All spike chaff components (e.g. glume, lemma, palea, and rachis) complete their differentiation before anthesis (AN stage). The *ppd-1* alleles in mutant lines differentially affected the duration of developmental stages pre- and post-anthesis: they extended all phases before anthesis, but shortened other phases that take place after anthesis, for example grain filling (Table 3). A shorter grain filling period (by 17% between AN and PM, Table 3) in the mutants resulted in shrivelled grains (Table 4), which counteracted any previous positive effects measured on carpel size at anthesis (Table 3).

To further investigate the effects of *Ppd-1* loci on spike architecture traits (e.g. grain number per spike, grain weight per spike, spike chaff components), we dissected spikes into spikelets and florets to quantify the effects on spikelet and floret organs (e.g. carpel, glume, lemma, palea). *Ppd-1a* alleles from NILs significantly increased the dry weight of F1, F2, F3, F4 florets, resulting in an overall rise in individual spikelet dry weight by 69% (Table 4). Similarly, NILs markedly improved individual grain weight for all floret positions: 41% for floret F1, 47% for floret F2, 74% for floret F3 and 264% for floret F4 (Table 4). Grain weight contributes over 80% of total floret weight, and will increase with longer grain filling periods, as observed in NILs (Table 3). By contrast, the shorter grain filling phase experienced by the *ppd-1* mutant lines led to a reduction in individual grain weight (Table 4). We also determined the effects of *Ppd-1* alleles on lemma and palea. Compared with WT, *ppd-1* mutants accumulated more dry weight in lemmas at all floret positions: F1 (34%), F2 (49%), F3 (74%), and F4 florets (110%). (Table 4). NILs had the lowest grain number per spike of all *Ppd-1* genotype combinations studied here, they also exhibited the heaviest grains per spike, because their high individual grain weight effectively compensated for their low grain number per spike (Table 4). The phenotypic variants of WT, NILs and KL validated the effects of *Ppd-1* allelic variants on spike architecture and developmental traits and display the connections between between spike architecture and development.

### Spatiotemporal Transcriptome During Loral Development Reveals the Effects of *Ppd-1* Loci on Gene Expression

Spike chaff components (e.g. lemma, glume, palea, rachis) are photosynthetically active and accumulate assimilates during the pre-anthesis phase. Grain number and size are greatly influenced by floret primordia initiation and development during pre-anthesis, and will reflect assimilate partitioning between green tissues and grains. Because spike architecture is closely linked to floret development, a better understanding of the factors that determine spike architecture is critical to discover the mechanisms underlying floret primordia initiation, development and degeneration.

To monitor the development and abortion process of all floret primordia within individual spikelets, we used transcriptome analysis of entire spikelets at six developmental stages (WA, GA, YA, TP, HD, and AN, Figure 2). Since the number of floret primordia peaks during the GA stage, and most abort by the AN stage, we collected spikelets starting at the WA stage (preceding the GA stage) to form a baseline before floret primordia numbers reach their maximum. We also collected samples at the YA, TP, and HD stages (before AN) to monitor the progress of floral development and abortion.

**Figure 2.**
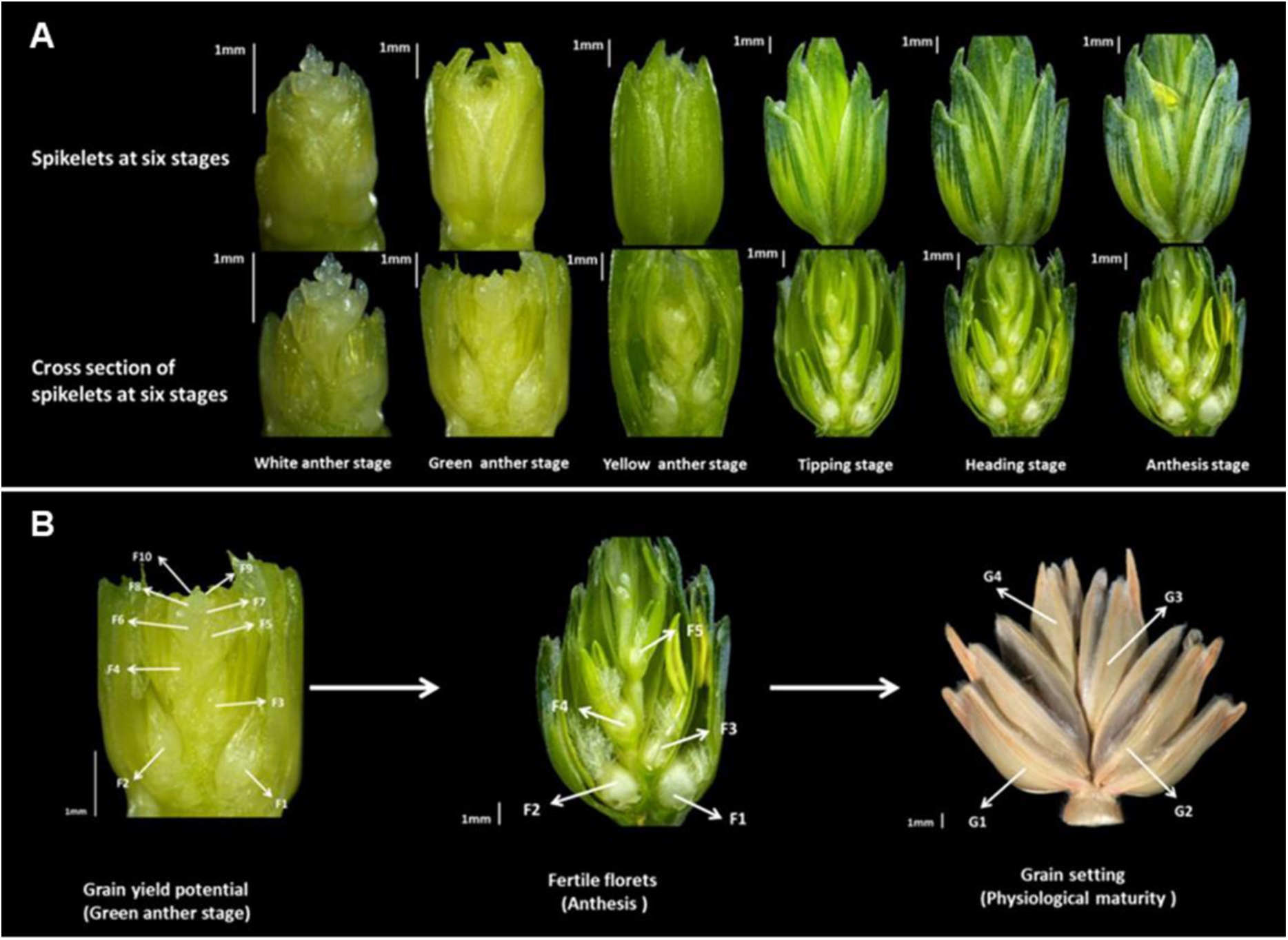
Definition of the developmental stages of wheat spikelets considered in this study. **(A)** Temporal dissection of spikelet development across six developmental stages. Top row: intact spikelets; bottom row: cross-sections through the central axis. Spikelets start forming floret organs (e.g. anther, carpel, lemma, palea) during the WA stage. Transcriptome data for the entire spikelet (without glumes) were collected at each stage, and transcriptome data for individual florets (florets 1, 2, 3, and 4-N: the floret primordia above the forth floret) were collected at the tipping, heading and anthesis stages. **(B)** Spatial dissection of spikelet development. The first three florets (F1, F2 and F3) generally set grains, but florets F4-N do not always, we collected samples for transcriptome analysis from all four positions. Florets F1, F2, F3, and F4-N develop into identifiable floret structures (e.g., anther, ovary) at the tipping stage (TS), we therefore started our dissection for transcriptome analysis at this stage. The number of floret primordia peaks at the green anther stage, and reflects grain yield potential. At anthesis, fewer than 50% of floret primordia have developed into fertile florets. At physiological maturity, fewer than 50% of floret primordia set grains (G1, G2, G3, and G4). Most floret primordia abort before anthesis.

To complement our spikelet transcriptome dataset, we also assessed individual florets at three stages (TP, HD, and AN). The basal florets within individual spikelets grow rapidly at these stages, while apical florets are already arrested or aborted. In hexaploid wheat, the first three basal florets (F1, F2, and F3, Figure 2) in spikelets from the central spike always set grains, while the fate of remaining florets (F4-N, Figure 2) is uncertain. We therefore determined the temporal gene expression profile of florets at three stages (TP, HD, and AN) and four positions (F1, F2, F3, and F4-N) within individual spikelets. Since we suspected that *Ppd-1* alleles would play a large role in shaping the transcriptome of spikelets and florets, we obtained samples from the three possible *Ppd-1* genotypes: wild type *Ppd-1* (WT; *Ppd-A1bB1bD1b*), one NIL (NIL; *Ppd-A1aB1aD1a*) and one mutant line (KL; *ppd-A1B1D1*).

The number of expressed genes detected in spikelets increased over our developmental time-course, ranging from 37,847 genes during WA to 40,524 during AN (Supplemental Data Set 1). To provide an overview of our expression data set, we generated a heatmap of all differentially-expressed genes (DEGs), which clearly showed waves of distinct gene expression pattern across stages and between genotypes (Figure 3A). We next identified genotype-specific genes involved in spikelet and floret development by comparing our transcriptome datasets across the three *Ppd-1* genotypes. The vast majority (85.1%) of DEGs displayed no preference for a *Ppd-1* genotype across all developmental stages. Genotype-specific genes related to floret development were by contrast relatively rare, accounting for only 1.9 to 3.6% of all DEGs (Figure 3B, Supplemental Data Set 1). Focusing on floret development, the three *Ppd-1* genotypes largely expressed the same gene sets, with an overlap of 82.8%. Again, genotype-specific differences in gene expression was rather minor, accounting only for 1.65 to 4% of DEGs (Supplemental Data Set 1).

**Figure 3.**
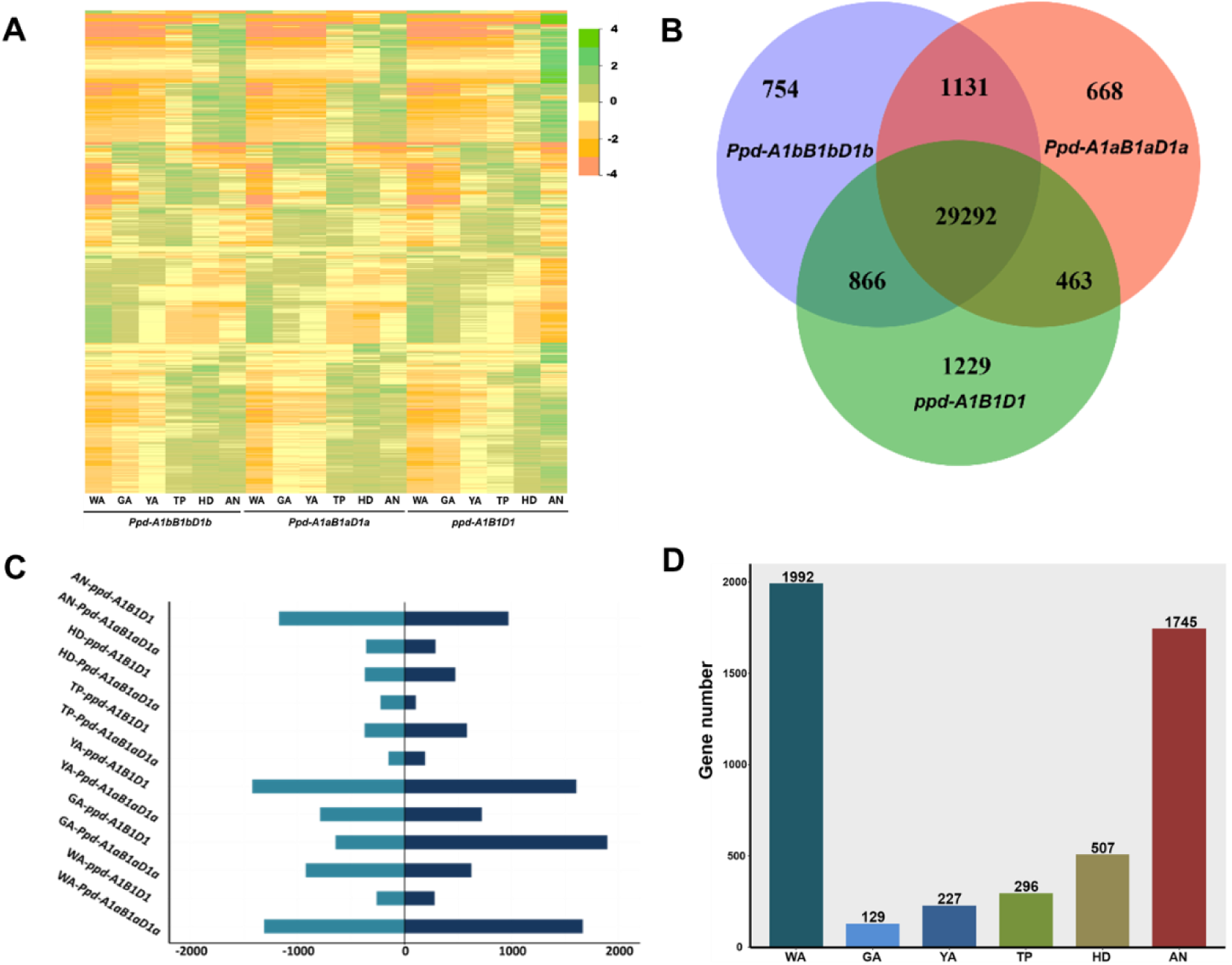
Expression profiles of spikelets across six developmental stages and three genotypes. **(A)** Heatmap of differentially expressed genes during spikelet development over the six developmental stages (WA, GA, YA, TP, HD, and AN) and in three genotypes (WT, NILs and mutant lines). Colors indicate relative expression (red, low expression; green, high expression). **(B)** Venn diagram of genes expressed in spikelets in each genotype. **(C)** Up- and down-regulated genes detected in NILs and mutant lines relative to WT. **(D)** Numbers of uniquely expressed genes specific for a single spikelet developmental stage. Four or five biological replicates were collected for each time point. Each replicate was a pool from at least three to ten plants.

During spikelet development, 9,399 genes were upregulated, against 8,012 downregulated genes over the course of the six stages (Figure 3C). When sorted by genotype, 3,593 genes were upregulated for all the six stages in the NIL when compared to WT. The *ppd-1* mutant line showed even more upregulated genes, with 5,806 genes relative to WT (Figure 3C). A comparable number of genes was downregulated relative to WT: 3,762 genes in the NIL and 4,250 genes in the mutant line (Figure 3C).

We then extracted stage-specific genes from our transcriptome across the six stages (WA, GA, YA, TP, HD, AN). Most genes expressed in spikelets overlapped with those expressed during each of the six stages, accounting for 87.9% of detected genes (40524 genes). Genes expressed during a single stage of spikelet development made up a much smaller fraction of expressed genes, with 1,992 genes only detected during WA and 1,745 genes specifically expressed during AN (the earliest and latest stages, respectively). Other developmental stages saw even smaller number of stage-specific genes: 129 genes during GA, 227 genes during YA, 296 genes during TP and 507 genes during HD (Figure 3D).

Finally, we investigated genes specifically expressed in florets across all three *Ppd-1* genotypes, three stages (TP, HD, and AN) and all four floret positions. Outside of genes whose expression was detected in all conditions, genotype-, stage- and position-specific genes comprised only 1.65–4%, 0.9–11.5% and 0.6–3.4% of all detected genes, respectively (Supplemental Data Set 1). When focusing on each genotype, we detected 1,568 genes that were expressed only in the NIL (*Ppd-A1a B1a D1a*), 977 in WT (*Ppd-A1b B1b D1b*) and 641 in the mutant line (*ppd-A1 B1 D1*).

### Spatiotemporal Changes in Functional Categories during Developmental Transition

To explore how the transcriptome landscape shapes spikelet and floret transitions during development, we investigated gene ontogeny (GO) terms over the course of our six development stages (Figure 2) and in the three *Ppd-1* genotypes. GO-term enrichment analysis during spikelet development revealed that terms such as “carbohydrate catabolic process”, “glucose metabolic process” and “response to hormone stimulus” were enriched early during spikelet development in all three genotypes (Figure 4A). The category “response to hormone stimulus” followed a similar spatial pattern during floret development within individual spikelets, as related GO terms were enriched only in basal florets within individual spikelets in all three genotypes (Supplemental Figure 1). These results corroborated previously published data demonstrating the critical importance of hormones and glucose in plant flower development (Song et al., 2018; Figueiredo et al., 2015; Béziat et al., 2017; Moore et al., 2003). These results also validate our own previous work that suggested that hormone- and sugar-related genes, such as the wheat homologues of *SUGAR SIGNALLING IN BARLEY 2, BRASSINOSTEROID INSENSITIVE 1* and *SUCROSE SYNTHASE 1*, were involved in floret development (Guo et al., 2017a).

**Figure 4.**
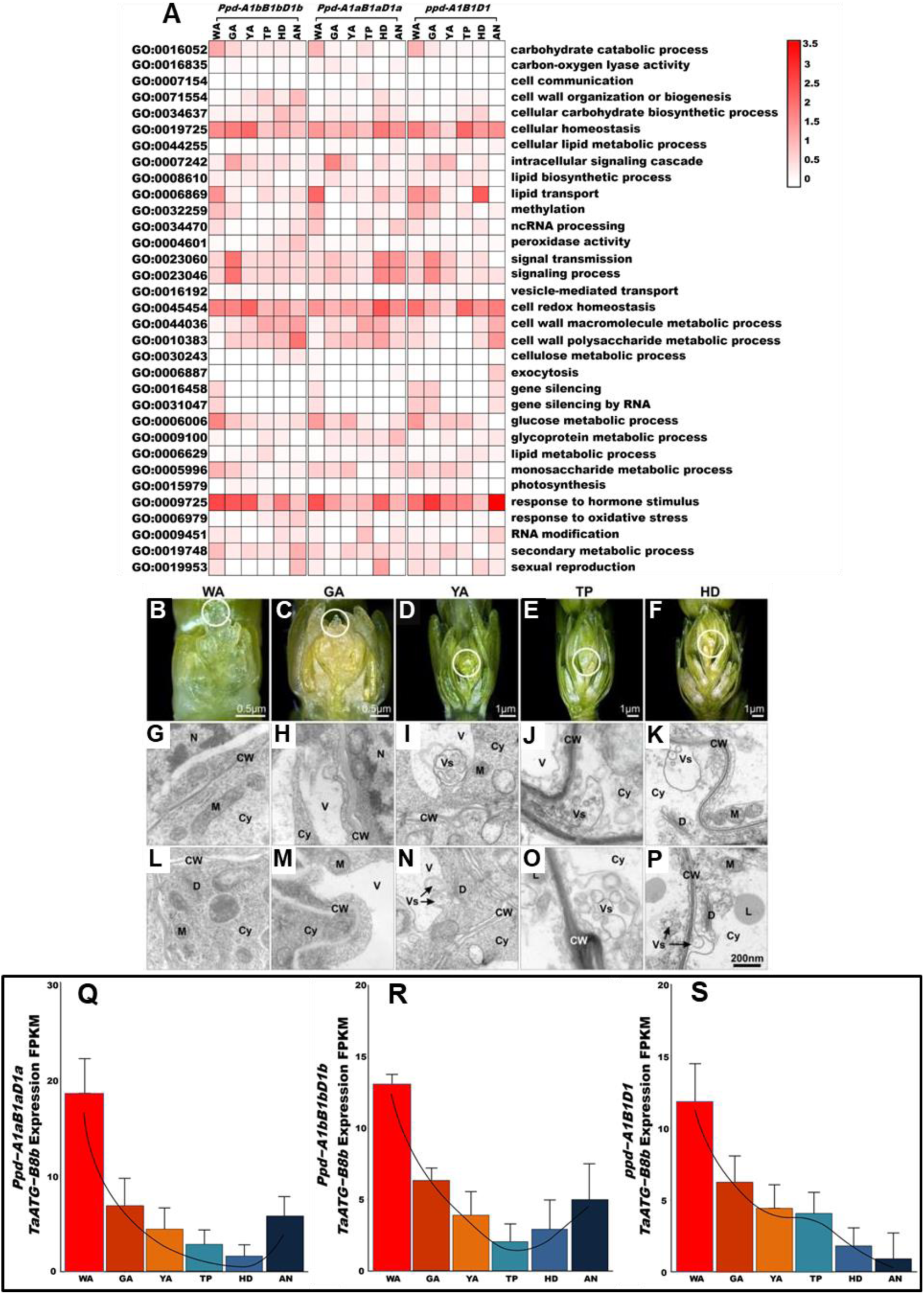
Temporal functional transition of spikelet gene and cellular morphology. **(A)** Enrichment in functional categories for genes expressed during spikelet development over six developmental stages in each of the three *Ppd-1* genotypes. The six stages (WA, GA, YA, TP, HD, and AN) are as defined in Figure 2. Red indicates dynamic changes across spikelet stages. **(B-P)** Modification of cellular morphology across six developmental stages. The paired figures (G-L, H-M, I-N, J-O, K-P) show the different positions of cell at the same stages. The transmission electron micrographs reveal the cellular morphology of the part indicated by the white circle. V, vacuole; Cy, cytoplasm; CW, cell wall; L, lipid drops; M, mitochondrion; **D, dictyosome;** Vs, vesicle. **(Q-S)** Expression levels of *TaATG-B8b* in the three *Ppd-1* genotypes (*Ppd-A1b B1b D1b, Ppd-A1a B1a D1a*, and *ppd-A1 B1 D1*) from our transcriptome datasets.

Conversely, terms such as “cell wall macromolecule metabolic process” and “cell wall polysaccharide metabolic process” were enriched at later spikelet developmental stages in all three genotypes. In support, we observed clear cellular modifications of apical florets as a function of their developmental stage (Figure 4B-4P). While no modifications can be seen during the WA stage, early in spikelet development (Figure 4G, 4L), the initiation of cellular modifications of apical florets is characterized by an increase in the size and number of vacuoles during the GA stage (Figure 4H, 4M
), followed by an increase in vesicle number at the YA stage (Figure 4I, 4N). Lipid droplets (L) become visible starting at the TP stage (Figure 4O). Cellular modifications occurred rapidly, starting at the GA stage, and almost all structures were lost by the HD stage: vesicles expanded to engulf much of the cell, lipid droplets took up a large fraction of cellular space, and the initial cell wall structure was lost (Figure 4J-4K, 4O-4P). Moreover, visible floral degeneration occurred after the GA stage, most obviously between the YA and HD stages (Figure 4D-4F; Table 2). Consistent with this observation, the GO terms “cellular homeostasis” and “cellular redox homeostasis” were significantly enriched among genes expressed at the YA, TP, and HD stages in all three *Ppd-1* genotypes. In this context, we observed more lipid droplets at the TP and HD stage (Figure 4P). However, we did not detect significant enrichment for GO term like “lipid biosynthetic” or “lipid metabolic process” at any of the stages we studied (Figure 4A, Supplemental Figure 1), which calls for further investigation in the future.

Autophagy may be involved in floral abortion in wheat (Ghiglione et al., 2008) and is closely associated with plant lipid homeostasis (Avin-Wittenberg et al., 2015; Fan et al., 2019). AUTOPHAGY-RELATED PROTEIN 8 (ATG8) plays a central role in the autophagy process (Behrends et al., 2010). We therefore determined the expression level of the wheat *ATG8* genes from the A and B genomes (D genome copy not expressed) to explore their potential role in spikelet development. Transcript levels of *TaATG-8bA* increased starting at the WA stage, while *TaATG-B8b* displayed the opposite pattern (Figure 4Q, 4R, 4S, Supplemental Data Set 2). In WT, within individual spikelet, *TaATG-A8b* expression decreased from basal (F1) to apical floret (F4) at TP and HD stages, but displayed an increased trend at AN stage (Supplemental Data Set 3). *TaATG-B8b* expression increased from basal (F1) to apical floret (F4), but displayed a decreased trend at AN stage (Supplemental Data Set 3). The ubiquitin-like autophagy-related protein ATG12 is required for the early steps of autophagy (Murrow et al., 2015). The expression of *TaATG-A12* was much higher than *TaATG-B12*, and was highly expressed at early developmental stages (WA, GA), concomitant with the first signs of floret primordia degeneration (Supplemental Data Set 2). Within individual spikelets, *TaATG-A12* expression increased from the basal floret (F1) to the apical floret (F4), which is consistent with the observed floret primordia phenotype: apical floret primordia are less advanced in their development at this stage, and will generally abort. KL (*ppd-A1B1D1*) decreased the expression of *TaATG-B8b* and *TaATG-A12* for spikelet at AN stage (Supplemental Data Set 2). Also, KL (*ppd-A1B1D1*) decreased the expression of *TaATG-B8b* for F1, F2, F3, F4 floret at TP stage (Supplemental Data Set 3). The expression of *Ppd-1* is very low in spikelet. Overall, our results indicate that *Ppd-1* loci may regulate floret abortion through their effects on the regulation of autophagy.

### Expression of PIFs MADS-box, SPL, EPFL and HD-ZIP Family Members

The transition to flowering is controlled by complex genetic networks. Members of multiple gene families, such as MADS-box, *SQUAMOSA PROMOTOR BINDING PROTEIN-LIKE* (*SPL*), *EPIDERMAL PATTERNING FACTOR-LIKE* (*EPFL*), *PHYTOCHROME INTERACTING FACTORs* (*PIFs*) and Homeodomain leucine zipper protein (*HD-ZIP*) transcription factors, play diverse roles in plant growth and development, including the guidance of inflorescence architecture and floral development. We therefore assessed transcript abundance for all 4 gene families in our transcriptome datasets. We detected 173 DEGs including genes in these four gene families and other genes related to spike development.

#### PHYTOCHROME INTERACTING FACTOR (PIF) Family

PHYTOCHROME INTERACTING FACTORS (PIFs) are basic helix-loop-helix transcription factors that interact physically with the red and far-red light photoreceptors phytochromes (Pham et al., 2018). Arabidopsis PIF1 is unique as it is the only PIF that exclusively represses seed germination in the dark (Oh et al., 2004). The *pif1* mutant *also* displays an early flowering phenotype (Wu et al., 2018). The wheat PIF1 homologue *TaPIF-D1* is one of the genes with highest expression (Figure 5A). The expression of *TaPIF-D1* decreased from WA to AN stages, but also responded to the status at the *Ppd-1* loci, as its expression decreased with later flowering times. The *ppd-1* mutant lines significantly lowered the expression of *TaPIF-D1* at the AN stage.

**Figure 5.**
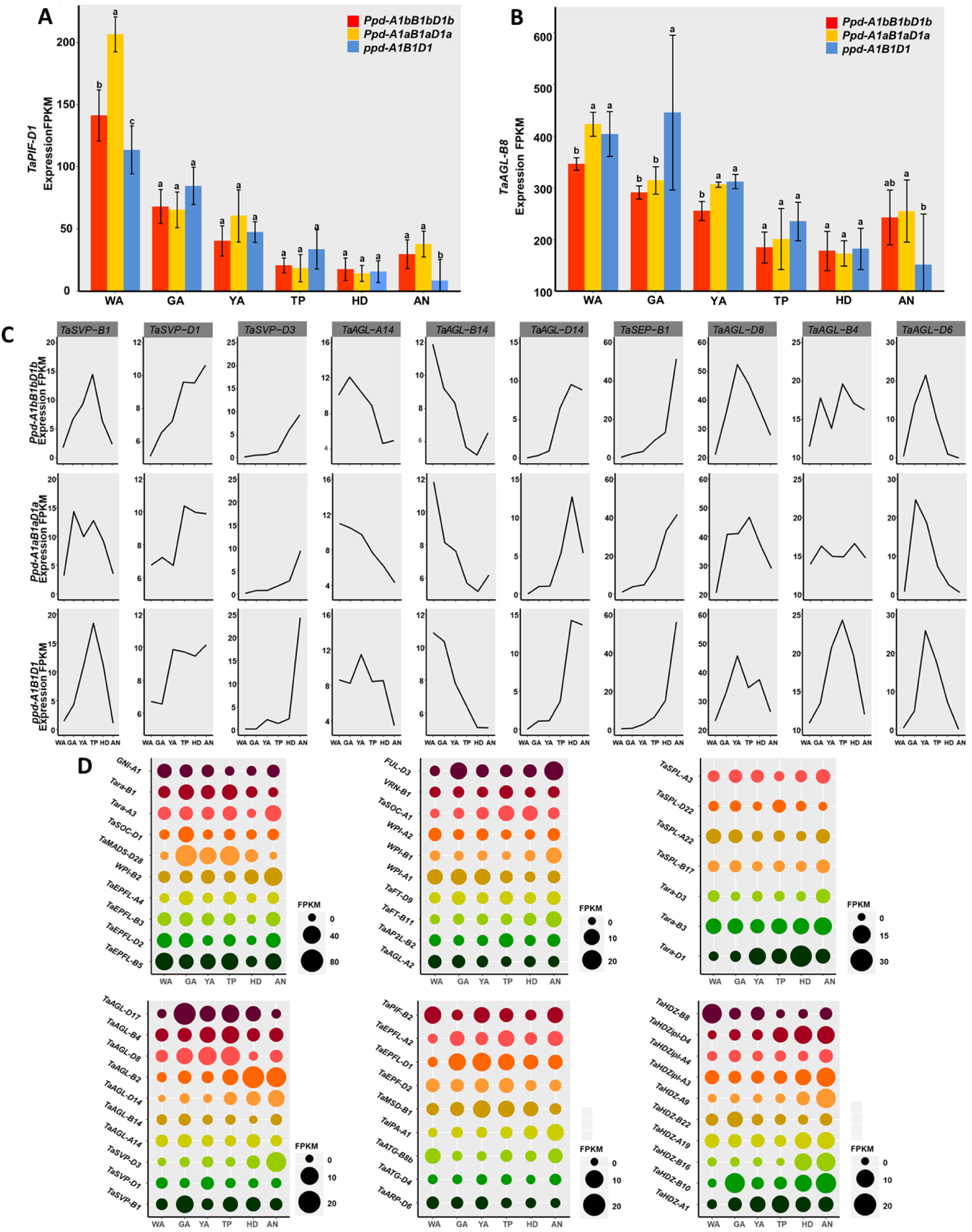
The expression dynamics of MADS-box family genes, *SQUAMOSA PROMOTOR BINDING PROTEIN-LIKE* (*SPL*) family, *EPIDERMAL PATTERNING FACTOR-LIKE* (*EPFL*) family, *PHYTOCHROME INTERACTING FACTOR* (*PIF*) family, *Homeodomain leucine zipper protein* (*HD-ZIP*) family and other genes regulating spikelet development. **(A-B)** the expression dynamics for the two genes (*TaPIF-D1, TaAGL-B8*) with highest expression levels. **(C)** the expression dynamics of genes in the three *Ppd-1* genotypes (*Ppd-A1bB1bD1b, Ppd-A1aB1aD1a*, and *ppd-A1B1D1*). **(D)** the differences between the highest and lowest expression levels between the three *Ppd-1* genotypes (*Ppd-A1bB1bD1b, Ppd-A1aB1aD1a*, and *ppd-A1B1D1*).

#### MADS-box Family

MADS-box genes specify floral organ identity (e.g. sepals, petals, stamens, and carpels). Members of the MADS-box transcription factor family have been studied in a wide range of species (Krizek and Fletcher, 2005), and their role in flower development is well understood at the molecular level. *SEPALLATA1* (also known as *AGAMOUS-LIKE2, AGL2*), *SEP2* (*AGL4*), *AGL6, AGL8* and *AGL14* are MADS-box genes with sequence similarity to the homeotic gene *AGAMOUS. AGL8* expression is not detected in flower primordia and accumulates in the developing carpels in Arabidopsis (Mandel and Yanofsky, 1995). *AGL8* also regulates the expression of genes required for cellular differentiation during flower development in Arabidopsis (Gu et al., 1998). We found that the expression of *TaAGL-B8* in wheat spikelets reached its highest expression level at the GA stage (446 FPKM) and then decreased during the HD and AN stages (100–200 FPKM), suggesting that *TaAGL-B8* may participate in carpel formation during the WA stage (Figure 5B, Supplemental Data Set 4). *ppd-1* mutant alleles significantly raised the expression of *TaAGL-B8* during the WA, GA, and YA stages, but later caused a drop in its expression at the AN stage (Supplemental Data Set 4).. Consistently, within spikelet, *TaAGL-B8* displays higher expression at apical florets (F4), which is less advanced florets (Supplemental Data Set 5). It suggests the important role of *TaAGL-B8* at early floral development. *Ppd-1* knockout significantly increases the expression of *TaAGL-B8* for spikelet at WA, GA and YA stages, but decrease its expression at AN stage.

In Arabidopsis, the *AGL2* transcript level remains high and constant throughout the floral meristem and in the primordia of all four floral organs: sepals, petals, stamens and carpels (Flanagan and Ma, 1994). We observed here that expression of *TaAGL-B2*/*TaSEP-B1* increased during wheat spikelet development, with lowest levels during the WA stage and reaching peak levels at the AN stage in all the three *Ppd-1* genotypes (Figure 5C, Supplemental Data Set 4). Consistently, we observed the higher expression of *TaAGL-A2* at apical florets (F4, less advanced floret) (Supplemental Data Set 5). It suggests the important role of *TaAGL2* at early stage, since the primordia are differentiating into floral organ of sepals, petals, stamens and carpels at early stage. *Ppd-1* knockout significantly increases the expression of *TaAGL-A2* for spikelet at AN stage (Figure 5D).

*AGL4* is also known as *SEP2.* The transcription factor FAR-RED ELONGATED HYPOCOTYL3 (FHY3) activates *AGL4*/*SEP2* expression to promote floral meristem determinacy (Li et al., 2016). By contrast, *TaAGL-B4* expression peaked at the TP stage (Figure 5C), which is consistent with the fast growth of floret/spikelet from between the TP and AN stages. A strong increase in *TaAGL-B4* expression in *ppd-1* mutant lines was observed at the TP stage.

*AGL6* expression is most abundant in developing flowers and ovules (Ma et al., 1991; Schauer et al., 2009). Loss of function of the rice and maize *AGL6* homologues affects floral differentiation (Ohmori et al., 2009; Thompson et al., 2009). The *APETALA2*-*like* gene *AP2L2* plays a critical role in specifying axillary floral meristems and lemma identity (Debernardi et al., 2019). *TaAGL-D17* and *TaAP2L-B2* expression peaked during the GA and YA stages (the key phase of floret primordia developing into organs), suggesting their important roles in floret differentiation (Figure 5C, Supplemental Data Set 4). The peak in *TaAGL-D17* and *TaAP2L-B2* expression differed depending on the status of the Ppd-1 loci: WT showed a peak during the GA stage, whereas both NILs (with only the insensitive alleles *Ppd-A1a B1a D1a*) and the mutant lines (with complete loss of function at all three loci lines: *ppd-A1 B1 D1*) delay their peaks to the YA stage. Within individual spikelet, we detected higher gene expression of at basal florets (more advanced florets).

XAANTAL2 (XAL2/AGL14) is critical for floral meristem maintenance and determinacy (Pérez-Ruiz et al., 2015). We observed a decrease in the expression of *TaAGL-A14* and *TaAGL-B14*, while the expression of *TaAGL-D14* increased, suggesting that each gene may carry out distinct functions during spikelet developmental (Figure 5C, Supplemental Data Set 4). Indeed, they reached peak expression levels at different stages: WA stage for *TaAGL-A14* and *TaAGL-B14*, and HD stage for *TaAGL-D14*, regardless of the genotype at the *Ppd-1* loci.

Wheat *PISTILLATA1* (*WPI1*) and *WPI2* encode the wheat homologues of the Arabidopsis *PISTILLATA* (*PI*) MADS-box gene. *WPI2* is the probable orthologue of the rice gene *OsMADS2* (Hama et al., 2004). The *WPI1* gene is expressed in the primordia of lodicules and stamens, but is not detected in the primordia of pistil-like stamens. The suppression of *OsMADS2* by RNA interference causes the flowers to fail to open and the transformation of lodicules into palea-like organs (Yadav et al., 2007; Prasad and Vijayraghavan, 2003). From the WA to the AN stage, *WPI-A1* globally decreased in expression (Supplemental Data Set 4), while *WPI-B2* followed the opposite pattern (Supplemental Data Set 4), indicating that *WPI-A1* and *WPI-B2* may play different roles during floret/spikelet development. Within individual spikelet, *WPI-A1* and *WPI-B2* are highly expressed at all the four floret positions. *ppd-1* mutant alleles significantly decreased the expression of *WPI-B2* during the HD and AN stages.

Previous work found that *ful3* mutants showed delayed flowering time and reduced stem elongation, arguing that *FUL3* plays a critical role in spikelet development (Li et al., 2019). Wheat *TaMADS28* is the homologue of rice *OsMADS31*, which is likely involved in salt tolerance during rice seed germination (Yu et al., 2018). We observed an increase in the expression of *FUL-D3* and *TaMADS-D28* from WA stage, peak expression at YA/TP stage (around 40 FPKM) and a later decrease from YA/TP to AN (Figure 5D, Supplemental Data Set 4). These results suggest that *FUL-D3* and *TaMADS-D28* may play a role at YA/TP stage, which corresponds to the spike booting phase in wheat (Guo et al., 2018). *ppd-1* mutant lines showed a delay in the peak of *TaMADS-D28* expression from the YA to the TP stage. Interestingly, we detected very low expression of *TaMADS-D28* at basal florets and much higher expression at apical florets. *Ppd-1* knockout delays the peak of *TaMADS-D28* expression from YA to TP stage and decreases its expression at apical florets. *FUL-D3* is highly expressed at apical florets (F3, F4) (Supplemental Data Set 5).

*FLOWERING LOCUS T* (*FT*) encodes a small globular protein that moves from the leaves to the shoot apex through the phloem (Wigge, 2011). At the shoot apical meristem (SAM), FT interacts with FD to promote the transition of the vegetative meristem into a reproductive inflorescence meristem (Wigge et al., 2005; Collani et al., 2019). In this study, *TaFT-B11* and *TaFT-D9* showed a consistent increasing trend from WA to AN stage in the wheat spike, and high expression in all the florets (Supplemental Data Set 4). *TaFT-B11* and *TaFT-D9* show a gradual increase in expression from the WA to the AN stage (Supplemental Data Set 4), suggesting a role during late floret/spikelet development. *ppd-1* mutant alleles significantly increased the peak of *TaFT-B11* expression, while decreasing the peak of *TaFT-D9* expression.

The MADS-box transcription factor FUL and SUPPRESSOR OF OVEREXPRESSION OF CONSTANS 1 (SOC1) redundantly promote flowering (Balanzà et al., 2014) by repressing the expression of *SHORT VEGETATIVE PHASE* (*SVP*). During early flower development, APETALA1 (AP1) represses the expression of two redundant flowering time genes, *SOC1* and SVP, to prevent floral reversion. During late flower development, such repression is necessary to activate *SEPALATA3* (*SEP3*) (Mutasa-Göttgens and Hedden, 2009). In our transcriptome dataset, *TaSOC-D1* expression decreased between the WA and TP/HD/AN stages (Supplemental Data Set 4), which would suggest a key role in early floral development. By contrast, *TaSVP-61* expression peaked later (during the GA/YA/TP/HD stages), while *TaSVP-D1* expression increased from WA to AN (Supplemental Data Set 4), indicating their variable roles in floral development. Also, we observed relatively high expression of *TaSOC-D1* at apical florets (F3, F4, less advanced florets) (Supplemental Data Set 5). *TaSVP-61* expression peaked during the GA stage in NILs (with photoperiod-insensitive *Ppd-1* alleles), while WT and mutant lines both delayed maximum expression to the TP stage.

In wheat, *VERNALIZATION1* (*VRN1*) regulates the transition from the vegetative to the reproductive phase, as well as the switch between winter and spring growth habits (Yan et al., 2003). The MADS-box genes *VRN1, FUL2* and *FUL3* all play critical and redundant roles in in the fate of the upper spikelet ridge and the suppression of the lower leaf ridge (Li et al., 2019). *VRN1* expression peaked at the YA/TP stage and dropped to its lowest levels during the AN stage (Supplemental Data Set 4), suggesting a specific role at the YA/TP stage. For individual floret, the highest expression of *VRN-B1 was* observed at apical florets (Supplemental Data Set 5). *ppd-1* mutant alleles delayed the peak of *VRN1* expression from YA to TP stage. Also, it decreases the expression at apical florets.

#### SQUAMOSA PROMOTOR BINDING PROTEIN-LIKE (SPL) Family

The decision to flower integrates multiple signals, one of them being plant age. This age-dependent pathway is controlled by the microRNA (miR) miR156 and its target genes *SQUAMOSA PROMOTER BINDING-LIKEs* (*SPLs*) (Cheng, 2004). SPLs are a key hub transcription factors for several flowering pathways in Arabidopsis (Hong and Jackson, 2015). It is unknown whether SPLs regulate flowering in wheat, or how. *SPL3* can mediate the activation of *APETALA1, LEAFY*, and *FRUITFULL* by the FT–FD complex to induce flowering under inductive long day photoperiods in Arabidopsis (Jung et al., 2016; Yamaguchi et al., 2009). *SEPALLATA3* (*SEP3*) expression was affected in response to altered *SPL3* and *FT* expression in the leaf and shoot apical regions in the regulation of flowering in Arabidopsis in response to ambient temperatures (Lee et al., 2012). SBP transcription factors therefore play critical and varied roles in plant development. As in Arabidopsis, *SPL3* and *SPL17* were putative targets for tae-miR156 (Xia et al., 2014). We observed a pronounced drop in *TaSPL-A3* expression from WA/GA/YA to HD/AN stages (Supplemental Data Set 4), while *TaSPL-B17* expression decreased from WA to TP/HD/AN stages (Supplemental Data Set 4). *ppd-1* loss of function alleles strongly lowered the expression of both *TaSPL-A3* and *TaSPL-B17* at the AN stage.

#### HOMEODOMAIN LEUCINE ZIPPER PROTEIN (HD-ZIP) Family

HD-Zip proteins are unique to plants, and contain a homeodomain closely linked to a leucine zipper motif involved in dimerization and DNA binding (Ariel et al., 2007). HD-Zip proteins are known to play crucial roles in plant development. The genes in HD-Zip family show variable roles at the different floret/spikelet development, and wheat HD-Zip genes are no exception. Transcript levels for some genes (e.g. *TaHDZ-A1, TaHDZ-B10, TaHDZipI-D4*) increased from low levels at the WA stage to reach their peak at TP/HD stages (Supplemental Data Set 4). Other genes (e.g. *TaHDZ-B22, TaHDZipI-D4*) saw a decrease in their expression level from WA, to reach their lowest level during late stages (Supplemental Data Set 4). *ppd-1* mutant alleles lowered the expression of *TaHDZipI-A3, TaHDZipI-A4, TaHDZipI-D4, TaHDZ-B10* and *TaHDZ-A19* during the AN stage, but raised expression levels of *TaHDZ-A9* and *TaHDZ-B16* at the same time. The *GRAIN NUMBER INCREASE 1* (*GNI1*) gene encodes an HD-Zip I transcription factor. The mutant *GNI-A1* gene is responsible for increased floret fertility in wheat (Sakuma et al., 2019). In our dataset, wild type *GNI-A1* expression clearly decreased from WA to YA (Supplemental Data Set 4), which is consistent with previous work (Sakuma et al., 2019). Within individual spikelet, wild type *GNI-A1* expression was mainly evident in the most apical Floret4-N position (Supplemental Data Set 5). It indicates the critical role in the determination of floret abortion beyond F3.

#### EPIDERMAL PATTERNING FACTOR-LIKE (EPFL) Family

Secreted peptides mediate intercellular communication (Tavormina et al., 2015). Several secreted peptides in the EPIDERMAL PATTERNING FACTOR-LIKE (EPFL) family regulate plant growth and development, including inflorescence and floret growth and development. *EPFL2* and *EPFL3* promote shoot growth and organ elongation through the regulation of plant hormones (Kosentka et al., 2019; Tameshige et al., 2016; Uchida et al., 2012). We identified a new *EPFL* gene and named it *TaEPFL-B5* (Supplemental Data Set 4). This new EPFL gene reached its highest transcript levels at the GA stage and its lowest transcript levels at the HD stage.

Genetic analysis in Arabidopsis demonstrated that four putative ligands (EPFL1, EPFL2, EPFL4, and EPFL6) function redundantly in the shoot apical meristem (SAM) by promoting organ elongation (Abrash et al., 2011; Uchida et al., 2012). The wheat gene *TaEPFL1* is required for wheat stamen development (Sun et al., 2019). In our dataset, *TaEPFL1* expression was high at the GA/YA stage (Supplemental Data Set 4), which is the main stage during which floral organ identity is established. We also saw that the expression of other wheat *EPFL* genes (*TaEPFL-D2, TaEPFL-B3, TaEPFL-A4, TaEPFL5* and *TaEPFL-B5*) all decreased between the WA and AN stages (Supplemental Data Set 4), suggesting that their roles may be restricted to the early stages of wheat spikelet development. *ppd-1* mutant alleles lowered transcript levels for *TaEPFL-D2, TaEPFL-B3, TaEPFL-A4, TaEPFL5* and *TaEPFL-B5* at the AN stage.

#### Additional Genes Related to Spike Development

In maize (*Zea mays*), *ramosa1* (*ra1*) encodes a C2H2 zinc-finger transcription factor, while *ra3* encodes a trehalose-phosphate phosphatase (Vollbrecht et al., 2005). Both RA1 and RA3 proteins colocalize in a narrow arc of cells at the base of the spikelet-pair meristem. *ra3* mutants reduce meristem determinacy and increase tassel branching (Eveland et al., 2014; Satoh-Nagasawa et al., 2006). The histone H2A variant H2A.Z is inserted into chromatin through the highly conserved SWR1 complex (Mizuguchi et al., 2004). Disrupting the activity of SWR1 by silencing one of its key conserved components, *ACTIN RELATED PROTEIN 6* (*ARP6*), triggers floret abortion in *Brachypodium distachyon* (Boden et al., 2013). The expression of the wheat genes *TaARP-D6, Tara-B1-like* and *Tara-A3* decreased from early stages (WA, GA) to the AN stage in our dataset (Supplemental Data Set 4), indicating that any role will be specific to early floret/spikelet development.

The rice transcription factor IDEAL PLANT ARCHITECTURE 1 (IPA1) reduces unproductive tillers and increases grains per panicle, which results in improved yield (Wang et al., 2018). We detected a clear raise in *TaIPA-A1* transcript levels between the WA and AN stages (Supplemental Data Set 4), suggesting a role in late floret/spikelet development.

In summary, we can distinguish sets of genes based on their peaks of expression differences along spikelet and floret developmental stages (Figure 5D). Genes for spikelet development with early peaks (WA and GA stages) include *TaMADS-D28, TaEPFL-B5, WPI-A1, TaAGL-D17*, and *TaHDZ-B8.* The next wave of genes (*TaSOC-A1, TaSVP-B1, TaHDZ-A1, TaAGL-D8, TaEPFL1-6D*, and *TaMSD1-2B*) reached their peak in transcript levels during the middle of the spike booting stages (YA, TP). Finally, late genes (*Tara-D1, Tara-A3, WPI-B2, FUL-D3, TaHDZipl-D4, TaHDZipl-A3, TaHD-A9, TaHDZ-B16, TaHDZ-B10, TaSEP-B1, TaSVP-D3* and *TaIPA-A1*) were maximally detected during late floret/spikelet developmental stages (HD, AN) (Figure 5D).

Genes for floret development with peaks of expression differences at floret F1 at different stages include *TaHDZ-A1*(F1-TP, F1-HD), *TaSEP-B1* (F1-AN), *TaHDZ-A1* (F1-TP, F1-HD), *TaEPFL-A2* (F1-TP, F1-HD), *and TaSOC-A1 (F1-TP).* The next wave of genes (*FUL-D3* (F2-TP, F2-HD), *TaAGL-D17* (F2-TP), *TaSVP-B1* (F2-TP), *TaSVP-D3* (F2-AN), *TaSOC-A1* (F2-HD), *Tara-D1* (F2-TP, F2-HD), *TaHDZ-B10* (F2-HD), and *Tara-B3* (F2-HD) reached their peaks in transcript levels at floret F2. The third wave of genes (*TaAGL-D17* (F3-HD), *TaEPFL-B10* (F3-AN), *TaAGL-D17* (F3-HD) reached their peaks of expression differences at floret F3. Finally, the largest differences of gene expression of *TaMADS-D28* (F4-TP, F4-HD) and *TaEPFL-B10* (F4-TP) were detected at floret F4 (Supplemental Figure 2 and Data Set 5). The results suggest that *Ppd-1* loci may regulate spike architecture through their effects on the regulation of floret development, which is based on the effects of the gene expression related to floret development.

## Discussion

One aim of our study was to determine the effects of various *Ppd-1* alleles on spike architecture and development, with the goal to identify new potential strategies for higher yield in wheat cultivars (Figure 6). We also catalogued the effects of *Ppd-1* alleles on gene expression patterns during spike, spikelet and floret development, which all contribute to spike architecture in wheat (Figure 6).

**Figure 6.**
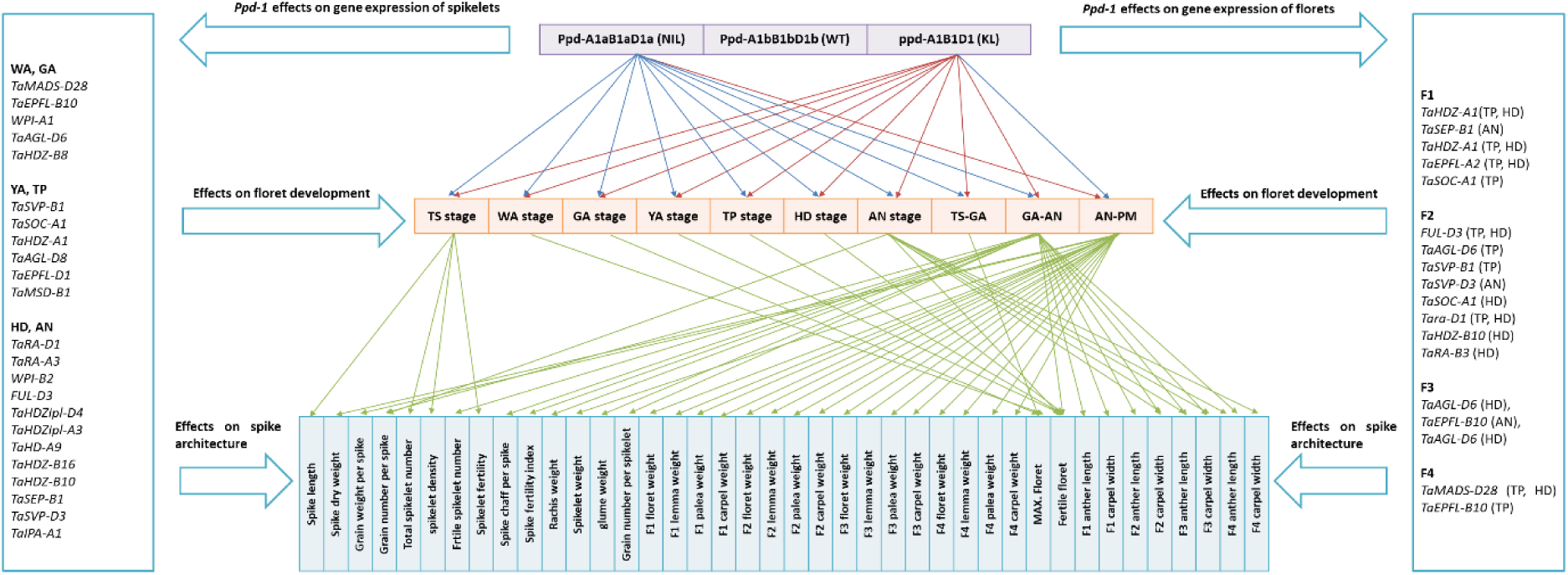
Connections between spike architecture and floral development traits. The blue lines and red lines indicate positive and negative effects, respectively. The green lines suggest the close connections between spike architecture and floret development traits.

We can summarize our study with five main conclusions: First, modulating the duration of spikelet primordia initiation did not always affect spikelet number. Near-Isogenic lines (NILs) carrying photoperiod-insensitive alleles at all three *Ppd-1* loci accelerated flowering time and thus decreased time to terminal spikelet and reduced spikelet number. However, delaying the time necessary to reach the terminal spikelet stage, as in *ppd-1* mutant lines did not significantly influence spikelet number. Previous work had shown that *FT1, FT2*, and *Earliness per se* (*Eps*) genes may increase spikelet number by extending the duration of spikelet primordia initiation (Lewis et al., 2008; Finnegan et al., 2018; Shaw et al., 2019). Here, our work suggests that increasing the duration of spikelet primordia initiation with loss of function *ppd-A1B1D1* alleles will not necessarily improve spikelet number. We conclude that *Ppd-1* loci have an intrinsic effect on spikelet number, which regulate the spikelet initiation. In other words, although *ppd-A1B1D1* genotypes may slow the rate of spikelet primordia, this effect is counter-balanced by an extension of spikelet primordia formation. Consistent with this hypothesis, *Ppd-1, Eps* and *FT1* genes were reported to influence spikelet number by altering the rate of spikelet initiation (Prieto et al., 2019; Dixon et al., 2018; Ochagavía et al., 2018). Further work will be needed to investigate the mechanism of the spikelet initiation in more details.

A second conclusion of our work is that extending the stem elongation phase (here by the use of the loss of function *ppd-A1 B1 D1* alleles) resulted in an improvement of spike chaff components (e.g. rachis, glume, lemma, palea). Again, the converse does not hold: the reduction in the stem elongation phase induced by NILs and their photoperiod-insensitive alleles (*Ppd-A1aB1aD1a*) did not lead to more floret organs (lemma, palea) at different positions, most likely due to the presence of other floral inhibitor genes e.g. *GNI-A1* (Sakuma et al., 2019). *Ppd-1* genes greatly affect the rate and duration of floret development (Pérez-Gianmarco et al., 2019; González et al., 2005), which may further regulate floret organs (e.g. lemma, palea). The vascular system within the rachis is critical for grain setting in wheat (Hanif and Langer, 1972). However, information about the effects of *Ppd-1* on rachis is scarce. We observed a marked effect of *Ppd-1* genes on rachis dry weight (Table 4). Any influence on the rachis vascular system may provide a potential explanation for the effects of *Ppd-1* on fertile florets and grain number by modulating the influx of assimilates into developing grains (Wolde and Schnurbusch, 2019).

A third conclusion stems from the observation that extending the pre-anthesis phase (as with the *ppd-1* triple mutants *ppd-A1B1D1*) paradoxically reduced phase duration of grain filling post-anthesis resulting in shrivelled grains. Similarly, extending the stem elongation phase caused carpel width increases that were reversed by a shortening of the duration of grain filling. These results indicate the existence of a fine balance between flowering time and grain number: prolonging flowering time may increase fertile floret number/grain number, but over-extension of the flowering window may reduce the duration of grain filling and thus negatively influence grain yield. Attempts to raise yield by altering flowering time alone will therefore need to be tread carefully.

Fourth, although most traits such as carpel width were influenced by the genotypes at the *Ppd-*1 loci, anther length was relatively stable in NILs (*Ppd-A1aB1aD1a*). Likewise, carpel width responded more strongly to a lengthening of floret development duration (from TS to AN stages) than it did to an acceleration of this developmental transition. Of interest is our observations documenting that *Ppd-1* greatly influenced the expression pattern of some genes, which may provide a potential explanation and future avenue of exploration.

Finally, NILs (*Ppd-A1aB1aD1a*) triggered floral degeneration. The allelic status of the *Ppd-1* loci clearly control the rate of spikelet/floret primordia development (Prieto et al., 2018; Ochagavía et al., 2018), further supporting an intrinsic role for *Ppd-1* in spike development. The intrinsic effect of *Ppd-1* may also explain its effect on floral degeneration.

These five findings highlight connections between spike architecture and floret developmental traits. We further displayed a high-resolution temporal landscape of the wheat spike transcriptome that provides new insights into floret primordia initiation, development and abortion, illustrating the value of these data sets for investigating diverse aspects of floral development and degeneration. The variable effects of *Ppd-1* alleles on the expression pattern of genes that belong to different families (PIFs, MADS-box, SPL, EPFL and HD-ZIP) at different time windows provide potential explanations for their effects on floret development, and thus spike architecture (Figure 6).

Taken together, our results have revealed connections between spike architecture and floret developmental traits by measuring 51 phenotypic traits in 197 wheat accessions and three *Ppd-1* genotypes. Our data set will facilitate the development of future hypotheses and will fuel research into the determinants of floret primordia initiation, development and abortion as well as spike architecture in wheat. Our results suggest that *Ppd-1* can remodel spike architecture by regulating floral development in wheat.

## Methods

### Plant Materials and Tissue Preparation

We grew hexaploid wild type (WT) wheat (*Triticum aestivum* L. cv. Paragon, a spring wheat carrying photoperiod-sensitive alleles in all three genomes: the triple sensitive allele combination *Ppd-A1bB1bD1b*), as well as plants from two near-isogenic lines (NILs, triple insensitive allele combination *Ppd-A1aB1aD1a*) and two independent mutant lines (KLs, triple knockout, *ppd-A1B1D1*) in a greenhouse for phenotypic measurements and transcriptome analysis. Insensitive alleles were derived from the genotypes GS-100 (*Ppd-A1a*), Chinese Spring (CS, *Ppd-B1a*), and Sonora 64 (*Ppd-B1a* and *Ppd-D1a*). Mutant alleles originated from the genotypes Paragon (Nor *ppd-A1* null), Paragon (211a *ppd-B1* null), Paragon (319c *ppd-B1* null) and Paragon (Nor *ppd-D1* null). In more detail, the two NILs are: NIL1 (GS-100 2A + CS 2B + Sonora 64 2D), NIL2 (GS-100 2A + Sonora 64 2B + Sonora 64 2D). The two mutant lines are: KL1 (Nor *ppd-A1* null + 211a *ppd-B1* null + Nor *ppd-D1* null), KL2 (Nor *ppd-A1* null + 319c *ppd-B1* null + Nor *ppd-D1* null). The generation of NILs and mutant lines was described previously (Shaw et al., 2013; Bentley et al., 2013). We used all 5 genotypes for phenotypic characterization, and 3 genotypes for transcriptome analysis: WT, NIL2 and KL2.

We selected 197 European hexaploid winter wheat cultivars from a previous genome-wide association study (210 cultivars) (Guo et al., 2017b). We genotyped photoperiod-sensitivity and photoperiod-insensitivity of photoperiod (*Ppd-D1*) (chromosome 2D) in this 197 cultivars. Of the 197 wheat cultivars, 165 cultivars are photoperiod-sensitive and 32 cultivars are photoperiod-in sensitive. Finally, we determined the effects of photoperiod-sensitivity and photoperiod-insensitivity of photoperiod (*Ppd-D1*) on the phenotypes (Guo et al., 2017b, 2018).

We sowed grains in 96-well trays and germinated them under greenhouse conditions (photoperiod, 16 h/8 h, light/dark; temperature, 20°C/16°C) for 14 d. We then vernalized seedlings at 4°C for 28 d. following which we transferred them to 15°C for one week to gradually acclimatize. Finally, we transplanted seedlings into pots, one per pot (9 × 9 × 9 cm). Supplemental light was provided and plants were irrigated when required.

We conducted temporal-spatial dissection to obtain overall information on floret development and abortion at six floral developmental stages of wheat: the white anther (WA) stage, green anther (GA) stage (maximum number of floret primordium), yellow anther (YA) stage, tipping (TP) stage, heading (HD) stage and anthesis (AN) stage. Details about the determination of different developmental stages can be found in (Guo et al., 2016; Guo and Schnurbusch, 2015). At each stage, the main shoots of three plants from each genotype were randomly selected to determine the number of floret primordia per spikelet. We took the entire spikelet (without the glume, from the spikelets in the centre of the spike) at each stage for temporal transcriptome analysis of spikelet (Figure 2). We took individual florets from four positions (F1, F2, F3 and F4-N, Figure 2) within individual spikelets (from spikelets in the centre of the spike) for temporal-spatial transcriptome analysis of floret (Figure 2). We collected four to five biological replicates for each time point, each replicate being a pool from at least three to ten plants. Total RNA was extracted using TRIzol reagent (Invitrogen).

### Read Mapping and Transcript Profiling

We downloaded the Chinese Spring reference genome from https://urgi.versailles.inra.fr/download/iwgsc/IWGSC_RefSeq_Assemblies/v1.0/. We assessed the quality of raw data using Fastqc (http://www.bioinformatics.babraham.ac.uk/projects/fastqc/). We then processed raw RNA-seq reads by first removing low-quality reads using trimmomatic (Bolger et al., 2014) and then mapped trimmed reads to the Chinese Spring reference genome using Hisat2 (Kim et al., 2015). After alignment, we normalized raw counts for each wheat gene to fragments per kilobase million mapped reads (FPKM). Gene expression counts were tabulated with HTseq-count (Anders et al., 2015). We determined significant differences in gene expression based on FPKM counts with the DESeq2 (Love et al., 2014) package in R. If the expression of one gene is below 1, we considered the gene was not expressed, which was further used to determine the stage-specific and position-specific genes.

### Gene Annotation and Functional Enrichment Analysis

We obtained gene models and annotations from the International Wheat Genome Sequencing Consortium database (https://urgi.versailles.inra.fr/download/iwgsc/IWGSC_RefSeq_Annotations/v1.1/). We obtained Gene Ontology (GO) terms for wheat genes from the ensembl plant database (http://plants.ensembl.org/index.html). We performed GO enrichment analysis with AgriGO (Du et al., 2010) and applied a hypergeometric test with rich factor correction (adjusted *P* value < 0.05).

### Transmission Electron Microscopy

For comparative ultrastructural analysis, wheat spikelets at different developmental stages were used for combined conventional and microwave-assisted tissue preparation. Aldehyde fixation, substitution and resin embedding were performed as defined in Supplemental Protocol (Supplemental Table 1). Sectioning and ultrastructure analysis were performed as described previously (Daghma et al., 2011).

## Supporting information

Supplemental Data Sets1-5

## Acknowledgements

We thank Dr Simon Griffiths and the Germplasm Resources Unit at John Innes Centre (Norwich, UK) for providing grains of the *Ppd-1* genotypes. Marion Roeder (Leibniz Institute of Plant Genetics and Crop Plant Research) and Martin Ganal (Trait Genetics GmbH) provided the 197 wheat accessions and the information of *Ppd-D1* allele in these accessions. We thank Dr. Martin Mascher (Leibniz Institute of Plant Genetics and Crop Plant Research), Yingyin Yao (China Agricultural University) and Jun Xiao (Institute of Genetics and Developmental Biology, Chinese Academy of Sciences) for suggestions about writing and data analysis. We thank Kirsten Hoffie and Marion Benecke for technical support in transmission electron microscopy. This work was supported by the Strategic Priority Research Program of Chinese Academy of Sciences, Grant No. XDA/B 00000000. Research in the Schnurbusch lab received financial support from the HEISENBERG Program of the German Research Foundation (DFG), grant no. SCHN 768/8-1 and the IPK core budget.

## Author contributions

Z.G. and T.S. designed and supervised the experiments. Z.G. performed the experiments and wrote the manuscript. Y.L. and L.Z. performed data analyses and prepared figures. M.M. conducted the analysis of cellular morphology. All authors viewed and edited the manuscript.

---

**Supplemental Table 1.**
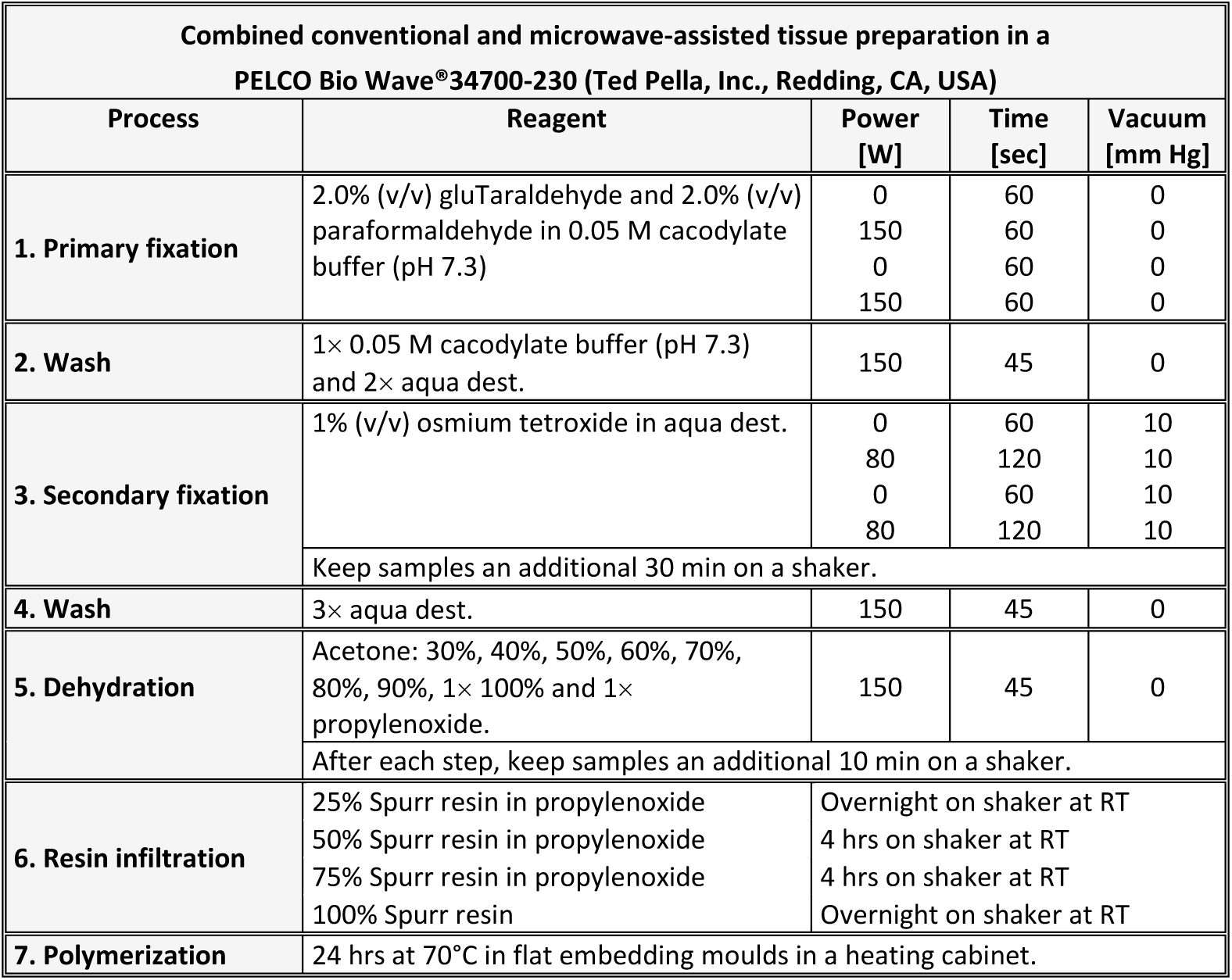
Protocol for combined conventional and microwave-assisted fixation, dehydration and Spurr resin embedding of wheat spikelets for ultrastructural analysis.

**Supplementary Figure 1.**
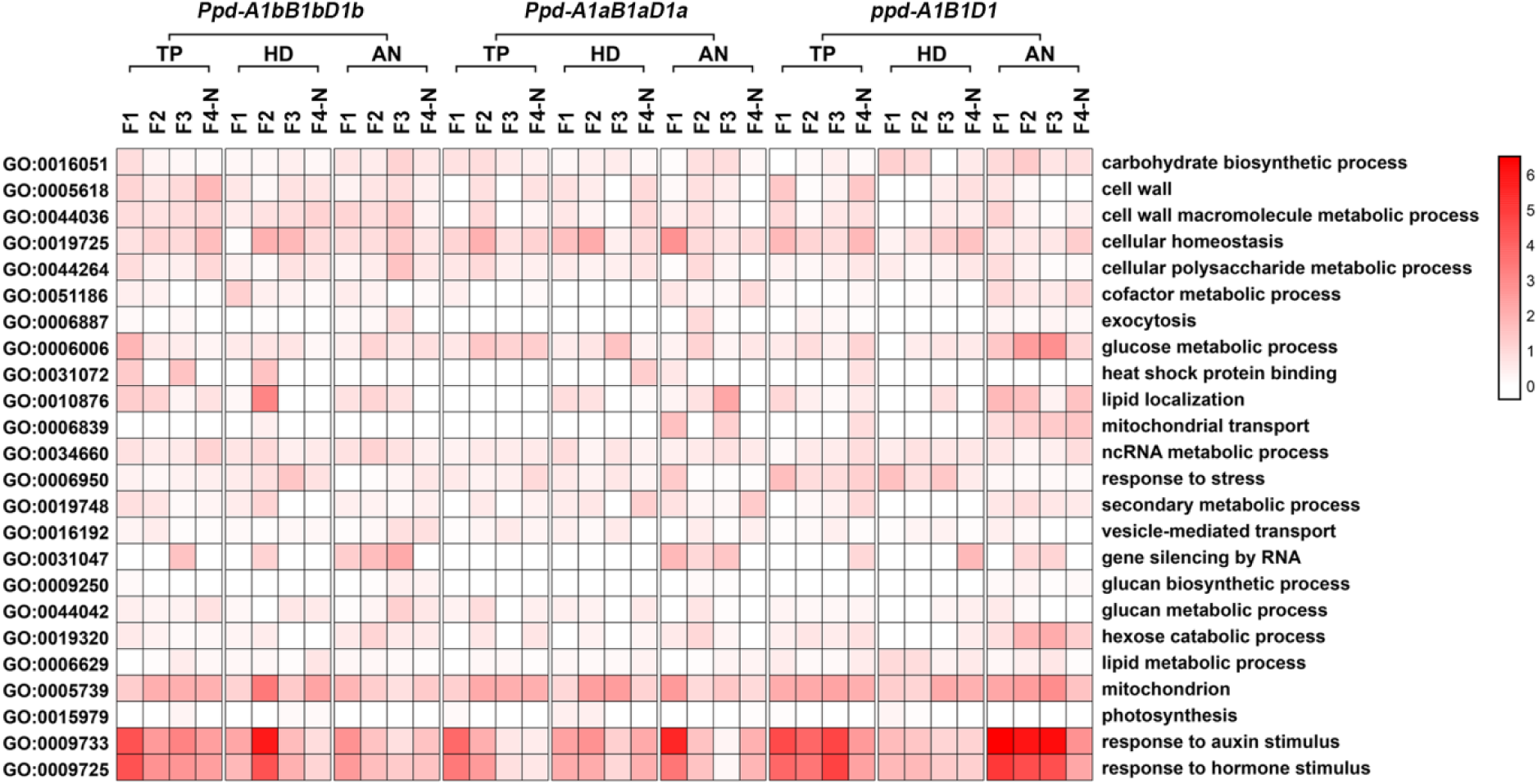
Temporal-spatial functional transition of floret genes across three developmental stages and four floret positions in the three Ppd-1 genotypes. The three stages (TP, HD and AN) and four floret positions (F2, F2, F3 and F4-N) are shown in Figure 1 and the three *Ppd-1* genotypes (*Ppd-A1bB1bD1b, Ppd-A1aB1aD1a*, and *ppd-A1B1D1*) in Table 1. The depth of the red colour indicates the dynamic changes of factors across the stages and positions.

**Supplemental Figure 2.**
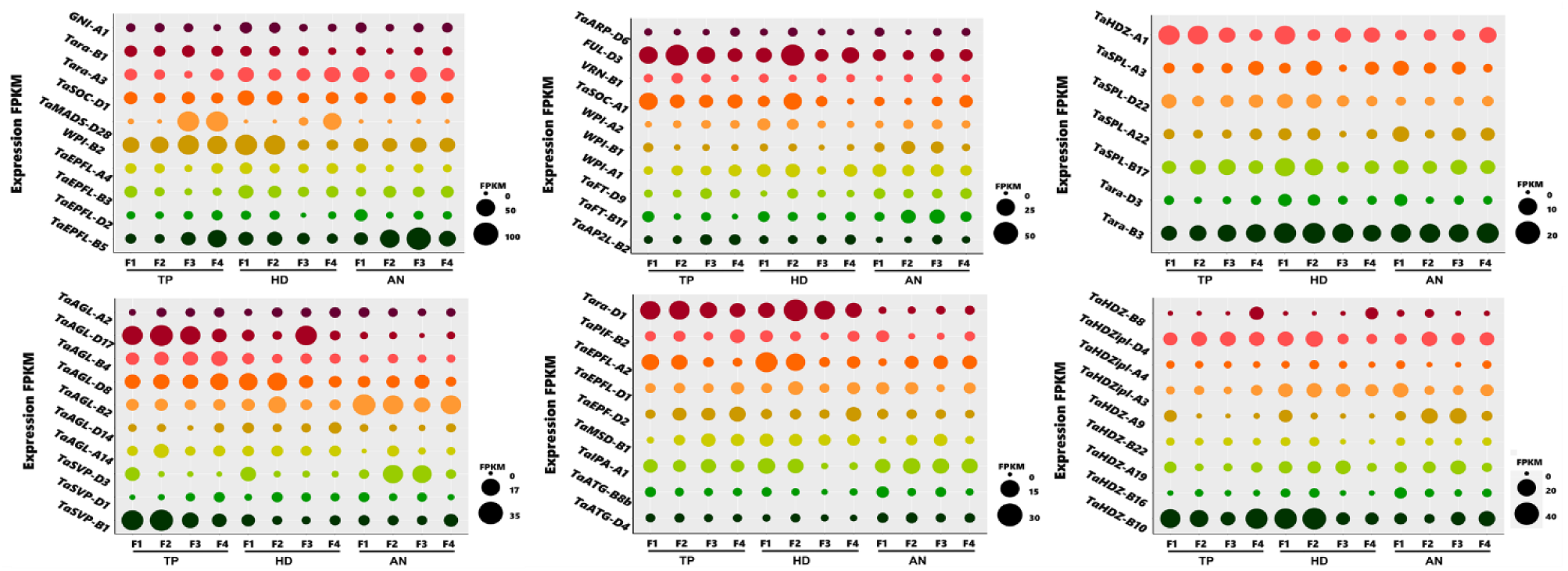
The differences between the highest and lowest expression levels between the three *Ppd-1* genotypes (*Ppd-A1bB1bD1b, Ppd-A1aB1aD1a*, and *ppd-A1B1D1*) for the MADS-box family genes, *SQUAMOSA PROMOTOR BINDING PROTEIN-LIKE* (*SPL*) family, *EPIDERMAL PATTERNING FACTOR-LIKE* (*EPFL*) family, *PHYTOCHROME INTERACTING FACTOR* (*PIF*) family, *Homeodomain leucine zipper protein* (*HD-ZIP*) family and other genes regulating floret development.

